# Climate and Morphology Drive Breeding Periods in Frogs

**DOI:** 10.1101/2022.07.21.501061

**Authors:** Bryan H Juarez, Lauren A O’Connell

## Abstract

**Aim:** Climate can have profound effects on reproductive behavior and physiology, especially in ectothermic animals. Breeding periods in amphibians have received little attention despite amphibian reliance on climate and water due to their reproductive biology and ecological diversity. The aim of this study is to determine how global climate impacts breeding periods in ectothermic animals through physiology, body size, and microhabitat.

**Location:** All continents, except Antarctica.

**Time period:** Breeding periods and climate both dating as far back as 1970.

**Major taxa studied:** 497 (7%) anuran species of 41 (76%) families.

**Methods:** We used phylogenetic comparative methods to analyze a global dataset of frog breeding periods, climate, body size, and microhabitat for 497 species.

**Results:** We found support for a global latitudinal gradient of breeding periods which are longer in the warmer, wetter tropics while shorter in the colder, dryer temperate zone. Latitudinal and non-latitudinal global patterns of breeding period were composites of the same patterns in the temperate and tropical zones. However, the effect of climate and body size in each zone is unique. Breeding periods displayed weak phylogenetic signal and were similar across microhabitats.

**Main conclusions:** Breeding periods show a global latitudinal gradient but this concept breaks down within the tropical zone. Our results are consistent with the importance of physiology in driving breeding periods and we describe how breeding period and body size may exhibit trade-offs which make latitudinal gradients context-dependent. Our results set within an ecophysiological framework have broad implications for understanding reproductive diversity in other ectothermic organisms.

## Introduction

Latitudinal gradients help reveal the origins of biodiversity. Examples include latitudinal gradients of species diversity (Pianka, 1966), geographical ranges (Stevens, 1989), niche breadth (Vázquez & Stevens 2004), biotic interactions (Schemske *et al*., 2009), and body size (Blackburn & Hawkins, 2004). Latitudinal gradients of species traits are important because they are evidence of evolution by natural selection in response to environmental differences. Behavioral traits responding to temporal changes in climate (phenology) are particularly helpful in understanding ecological adaptation (Wcislo, 1989; Blomberg *et al*., 2003). The diversification of behavior is often studied by measuring trait differences between one or more species exposed to various conditions in controlled environments or by determining how behavior is related to climate among many species through comparative approaches (Harvey & Purvis, 1991). This includes research on the effects of latitudinal climate on flowering and fruiting in plants (Badeck *et al*., 2004; Butt *et al*., 2015; Daru *et al*., 2019), hatching, emergence, and flight in arthropods (Kaspari *et al*., 2001; Śniegula *et al*., 2016), and migration, breeding, and biotic interactions in animals (reviewed in Cohen *et al*., 2018). Much of this research has benefited from using ectothermic organisms although research on amphibians is relatively limited. Since many amphibian groups are threatened by population declines, understanding how behavioral traits covary with climate fills a gap in our general understanding of phenology and informs future conservation strategies.

Frogs (Anura) are an ecologically diverse group (>7,400 species; AmphibiaWeb, 2023) ideal for studying phenological patterns. In addition to being ectothermic, reproductive cycles and calling behaviors in frogs vary with temperature, precipitation, and latitude (Wells, 2007). Additionally, anurans exhibit clear and rapid responses to climate. For example, warmer winters result in earlier breeding in frogs (Beattie, 1985; Reading, 2003; Neveu, 2009; Todd *et al*., 2011; While & Uller, 2014; Benard, 2015). However, relative to breeding start dates, little is known about how climate impacts breeding periods despite the stunning contrast of tropical species that breed year-round to explosive-breeding species that limit reproduction to only a few days of the year (Wells, 1977). Natural selection likely acts on breeding periods because longer periods of reproduction are associated with the production of more egg clutches (Bull & Shine, 1979; Morrison & Hero, 2003) and warm and wet climates (Morrison & Hero, 2003; Wells, 2007). One study on anurans found breeding durations (at the family level) were shorter in the dryer parts of Brazil (Forti *et al*., 2022). To our knowledge, no studies have examined climate effects on frog breeding phenology specifically at a broad taxonomic and global scale.

There are several non-exclusive hypotheses relating climate and ecology to frog breeding periods. For example, breeding periods could be constrained by cold and dry climates since frogs generally rely on warmth and water for reproduction. This ‘climate hypothesis’ predicts that species at lower latitudes should have longer breeding periods. Additionally, reproductive behaviors could vary across latitudes due to climatic differences between regions. This ‘zone hypothesis’ predicts that phenology may be influenced more by temperature in the temperate region and precipitation in the tropics (Cohen *et al*., 2018). Finally, extreme habitat differences may drive reproductive behaviors, as burrowing frogs living in deserts often have breeding lasting from only a few days (Wells, 1977). This ‘microhabitat hypothesis’ predicts that desert environments constrains breeding compared to other microhabitats.

In addition to the climate and ecology, body size may also be an important factor influencing phenology (Cohen *et al*., 2018). Larger body sizes in frogs are associated with larger clutches (Gomez-Mestre *et al*., 2012; Furness *et al*., 2022) and possibly access to larger-bodied prey (Wilson, 1975) or greater storage capacity of energy from food (as in mammals; Demment & Van Soest, 1985). The pace of life framework (Hille & Cooper, 2015) connects environmental variation with life history and body size. For example, it describes how warmer, wetter, and less seasonal environments (where growth and reproduction are not environmentally constrained) might select for larger body sizes, slower developmental rates, larger clutches, and many reproductive events (iteroparity). The converse applies to smaller species under the pace of life framework. Thus, we predict relatively larger body sizes are associated with longer breeding periods in warmer, wetter, and less seasonal environments and the converse for smaller species. We refer to these dual predictions as the ‘climate-size hypothesis’.

The goal of this study is to use the comparative approach to understand the climatic and morphological factors which might influence breeding periods, i.e. breeding durations at the population level (Wells, 1977), in frogs. Specifically, we test predictions from the climate, zone, and climate-size hypotheses to determine: 1) whether breeding periods are driven by latitude, geographical zone, or phylogenetic history, 2) whether longer breeding periods are associated with warm, wet, and less seasonal climates, 3) whether burrowing lifestyles result in shorter breeding periods, and 4) whether the relationship between breeding period and climate depends on body size.

## Material and Methods

### Coordinate and climate data

We obtained coordinate data for 497 species representing 41 of 54 anuran families from GBIF.org (GBIF.org, 2021a,b,c,d, accessed 3 October 2021). We describe our data cleaning procedures in Appendix S1 in Supporting Information. We subsetted the coordinate data to match the set of species for which we obtained behavior, phylogenetic, microhabitat, and body size data. This resulted in 848,572 coordinates (mean = 1,707 and range = 3–83,065 observations per species) from which we obtained species mean estimates of absolute latitude and climate data using WorldClim 2.1 (Fick & Hijmans, 2017), and envirem (Title & Bemmels, 2018) at 30 seconds (∼1km^2^) resolution. We used an equal-area Mollweide projection to avoid biases in land area. From these projections, we extracted from each coordinate variables affecting temperature and water availability covering all possible breeding period values (0–365 days). We reduced the effects of spatial autocorrelation by avoiding repeated sampling of the same (1km^2^) pixels. Specifically, this dataset included annual mean temperature, temperature seasonality, annual precipitation, precipitation seasonality, annual potential evapotranspiration (PET), PET seasonality, and topographic wetness. Topographic wetness is the land’s ability to retain water and is obtained from estimates of upslope incurrent areas and local slope (Sørensen *et al*., 2005). We calculated species means for each climate variable to analyze the dataset in a phylogenetic comparative framework.

### Breeding period data

We compiled species means of breeding period (sensu Wells, 1977) using data from the primary literature, AmphibiaWeb (2022), and online databases. We estimated the breeding period as the difference in calendar days between the breeding start and end dates, assuming 365 days, when the breeding period was not already reported in days. We then averaged these values across populations (N = 1–6, mean = 1) to obtain species means. Some records were reported as qualitative data (e.g., “early May”). When breeding periods were reported as starting or ending on a specific month, we ascribed values of either the first or last day of the month, respectively. When breeding periods were reported as the “early” or “late” parts of a month, we matched this to the end of the 1^st^ week and 3^rd^ week, respectively. Rarely, breeding periods were reported as specific seasons (e.g., “Summer” or “wet season”) and we defined these as the number of days corresponding to each season and country. Since explosive breeding is <30 days (Wells, 1977) and because we did not find any records of explosive breeding which took longer than 2 weeks, we conservatively assigned a period of 2 weeks to species whose only description of the breeding period was “explosive breeding”.

### Phylogenetic, habitat, and body size data

We obtained the consensus tree and 1,000 trees randomly sampled from the pseudoposterior distribution of trees from Jetz and Pyron (2018) to account for shared evolutionary history and phylogenetic uncertainty in our analyses. Briefly, Jetz and Pyron (2018) used PASTIS (Thomas *et al*., 2013) which combined taxonomic information from AmphibiaWeb and a molecular supermatrix of 5 mitochondrial and 10 nuclear genes (Pyron, 2014; Jetz & Pyron, 2018) to estimate a posterior distribution of trees. We collected microhabitat and maximum male body sizes (snout-vent length) from the primary literature and other datasets (Moen & Wiens, 2017; Womack & Bell, 2020; AmphibiaWeb, 2022; IUCN, 2022). Microhabitat categories included aquatic, arboreal, burrowing, terrestrial, torrential, semi-arboreal, semi-aquatic, and semi-burrowing (Moen & Wiens, 2017). A list of the sources cited in data tables is found in Appendix 1.

### Statistical analysis and visualization

We inspected the latitude and climate data in many ways to understand their relationships in a biogeographical context. All analyses and visualizations were done in R v4.2.1 (R Core Team, 2022). First, we plotted species means for each climate variable against absolute mean species latitude to characterize the latitudinal climate gradient in our data. Next, we centered and scaled each climate variable to perform principal component analysis (PCA) and understand the covariances among climate variables across species distributions. We also visualized spatial variation of breeding periods and climate data by estimating per-pixel mean breeding period and principal component scores at 100 km^2^ resolution using letsR v.4.0 (Vilela & Villalobos, 2015).

To understand how breeding periods and absolute latitudes are distributed across the phylogeny of frogs, we plotted average species breeding periods and absolute latitude along the tips of the consensus tree of Jetz and Pyron (2018) using *ggtree* v3.4.0 (Yu *et al*., 2017; Yu *et al*., 2018; Yu, 2020). To determine the extent to which phylogenetic history might explain our data, we estimated the phylogenetic signal (Adams, 2014) of breeding periods and absolute latitude and using *geomorph* v4.0.1 (Adams *et al*., 2021; Baken *et al*., 2021b). Additionally, we also obtained summary statistics of breeding periods and absolute latitudes for the four largest families represented in our dataset: Hylidae (N = 98), Myobatrachidae (N = 86), Bufonidae (N = 62), and Ranidae (N = 44). To test for a latitudinal gradient, we regressed breeding periods onto absolute latitude using phylogenetic generalized least squares (PGLS; Grafen, 1989; Martins & Hansen, 1997). PGLS regression uses a phylogenetic covariance matrix to account for residual correlation due to shared evolutionary history. Furthermore, we implemented PGLS regression through *RRPP* v1.3.1 (Collyer & Adams, 2018, 2019) which uses a residual randomized permutation procedure to fit linear models, evaluate significance, and calculate empirical effect sizes for model terms in all models. We used 9,999 iterations to evaluate significance in all models.

We tested whether colder, dryer, and more seasonal environments are associated with shorter breeding periods using principal component (PC) regression. PC regression avoids collinearity issues while helping us understand how breeding periods are linked to environments defined by covariation among climate variables (PC loadings). We combined principal component regression with PGLS regression by regressing species means of breeding period onto all seven principal components. We then compared the fit of this model to the fit of a phylogenetic ANCOVA model including body size, microhabitat, and interactions between body size and each principal component to determine how the relationship between breeding period and climate might depend on body size. We also performed pairwise comparisons of breeding periods among microhabitat categories to determine whether burrowing lifestyles are associated with shorter breeding periods. To account for multiple comparisons, we controlled Type I error rates (a = 0.05) using the Sidak-like step-down procedure in *mutoss* v0.1-12 (MuToss Coding Team *et al*., 2017). This method is more powerful than both the Sidak and Bonferroni methods (Holland & Copenhaver, 1987). We used the natural logarithm of body size to account for evolutionary allometry of body size. To test the zone hypothesis, we repeated the same modeling procedures above for two subsets containing either temperate or tropical zone species.

Finally, we evaluated whether the phylogenetic ANCOVA results above were robust to phylogenetic uncertainty. Following Baken *et al*. (2021a), we fit new PGLS models using each of the 1,000 trees from Jetz and Pyron (2018) to obtain *Z*-scores for each independent variable. This procedure resulted in 1,000 *Z*-scores for each model term from which we calculated the proportion of trees for which the effect size was significant corresponding to *Z* > 1.645 for empirically generated *Z*-scores (Collyer & Adams, 2013; Collyer *et al*., 2015).

## Results

### Climate data showed latitudinal and non-latitudinal differences

Relationships between species means of each climate variable and absolute latitude (Fig. 1) showed climate differences across the tropical and temperate zones. Overall, frogs in lower (tropical) latitudes live in warmer, wetter, and less seasonal areas, relative to frogs at higher latitudes. One important exception is that tropical frogs experience the most precipitation seasonality. Precipitation seasonality also increases with latitude in tropical frog habitats. Additionally, annual precipitation and precipitation seasonality is more variable than annual mean temperature and temperature seasonality among tropical frogs. Absolute latitude was highly correlated (*r* = −0.94; Fig. 2) with PC1 (52% variance; Table 1). However, 48% variance (PC’s 2–7) was independent of latitude (PC1).

**Figure 1.**
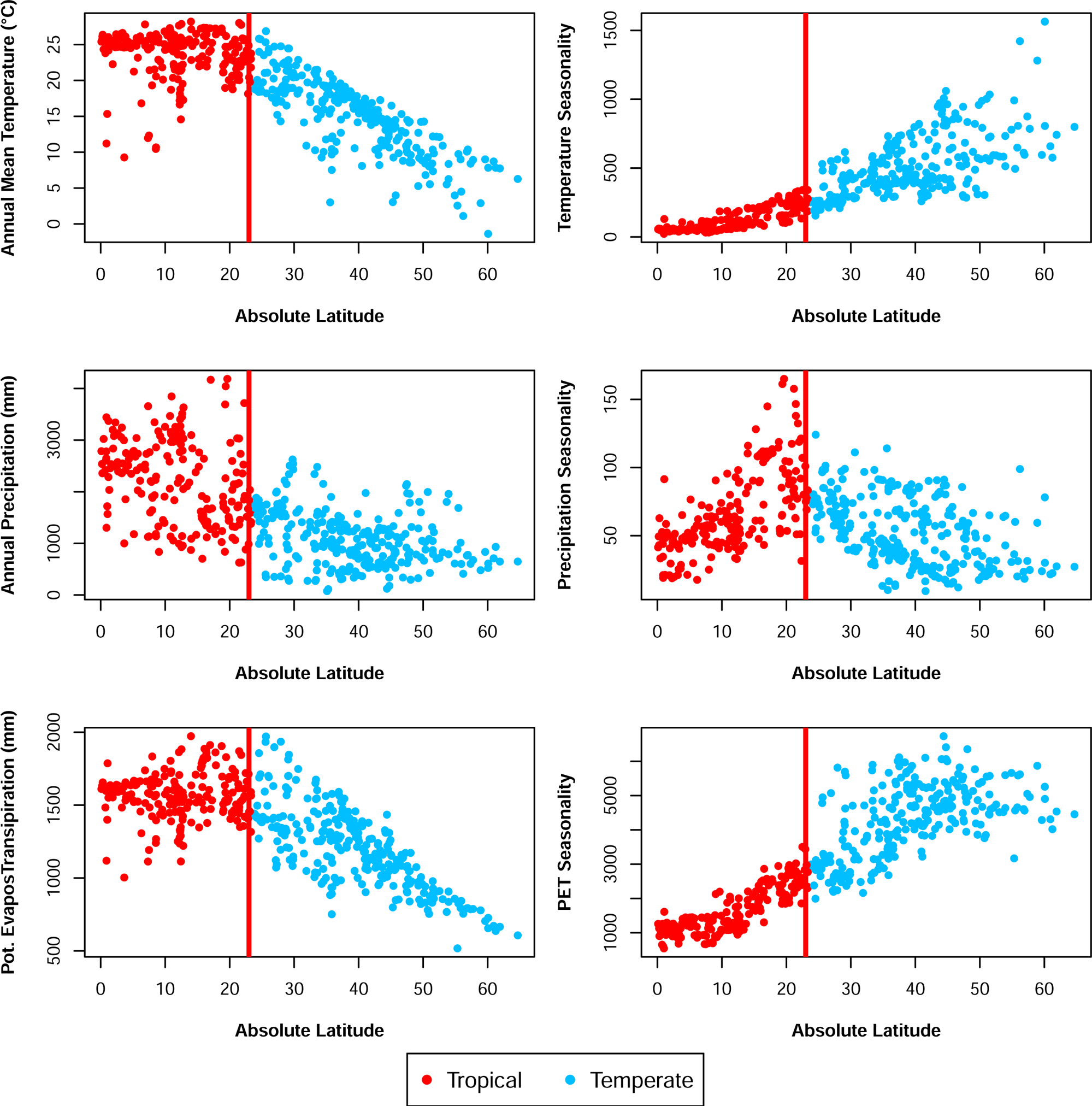
Relationships between species means climate and absolute latitude. Generally, tropical and temperate climates differ substantially relative to absolute latitude when considering annual mean temperature, temperature seasonality, annual precipitation, precipitation seasonality, potential evapotranspiration, and potential evapotranspiration seasonality. Red points indicate tropical species and blue points indicate temperate zone species. Vertical red line corresponds to 23.43628 decimal degrees.

**Figure 2.**
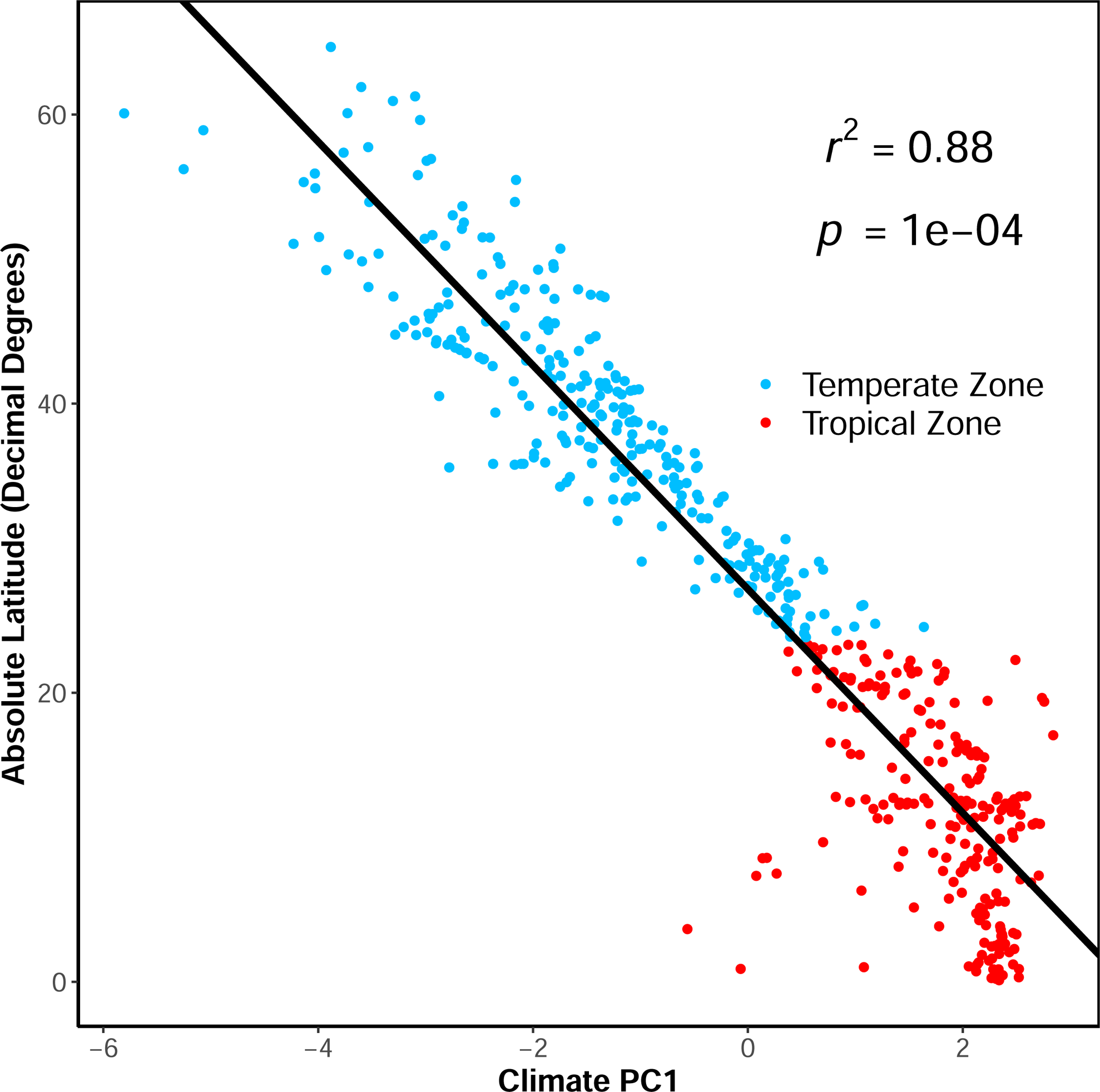
Relationship between species means of absolute latitude and Climate PC1. Blue points correspond to temperate zone species and red points are tropical zone species. Tropical and temperate zone cutoff used is 23.43628 decimal degrees. Absolute latitude is strongly and negatively correlated to PC1. *r^2^* = 0.88 and *p* < 0.0001.

**Table 1.** PCA summary and loadings of climate variables. PCA based on means for 497 species. Cumulative proportion of variance is given in parentheses next to PC heading. Loadings with a magnitude >=0.30 are in bold.

| Variable | PC1<br>(52) | PC2<br>(76) | PC3<br>(90) | PC4<br>(95) | PC5<br>(97) | PC6<br>(99) | PC7<br>(100) |
| --- | --- | --- | --- | --- | --- | --- | --- |
| Annual Mean Temperature | **0.47** | -0.27 | 0.08 | -0.01 | 0.25 | 0.16 | **0.78** |
| Temperature Seasonality | **-0.47** | -0.14 | -0.09 | **0.46** | 0.22 | **0.70** | 0.04 |
| **Annual Precipitation** | **0.38** | **0.38** | 0.13 | **0.75** | 0.25 | -0.21 | -0.14 |
| **Precipitation Seasonality** | 0.15 | -0.26 | **-0.90** | 0.22 | -0.17 | -0.16 | 0.00 |
| **Annual Potential Evapotranspiration (PET)** | **0.42** | **-0.40** | 0.01 | -0.21 | **0.47** | 0.23 | **-0.59** |
| **PET Seasonality** | **-0.45** | -0.28 | 0.01 | 0.03 | **0.60** | **-0.59** | 0.10 |
| **Topographic Wetness** | 0.01 | **-0.67** | **0.40** | **0.36** | **-0.48** | -0.16 | -0.10 |

### Breeding periods exhibit a latitudinal gradient

We examined phylogenetic patterns of variation and covariation for mean breeding period (mean = 148 days, range = 1–365 days, std. error = 4 days) and absolute latitude (mean = 27.17° N, range = 0.11–64.67° N, std. error = 0.71°). Each trait exhibited a weak phylogenetic signal, including breeding period (*K* = 0.19, *p* = 0.0001; Fig. 3) and absolute latitude (*K* = 0.51, *p* = 0.0001). Taxon-specific averages of breeding periods and absolute latitude matched the prediction of finding longer breeding periods at lower latitudes: hylid treefrogs (1.90° N) bred for 187 days, bufonid toads (18.24° N) bred for 114 days, and ranid water frogs (43.88° N) bred for 110 days. However, Australian and southeast Asian myobatrachid frogs bred for 143 days despite living in the temperate zone (33.85° S). Summary statistics suggested tropical species bred for longer periods, on average (mean = 174 days, range = 5–365 days, std. error = 7 days), relative to temperate species (mean = 128 days, range = 1–365 days, std. error = 5 days). In fact, we found longer breeding periods were weakly associated with lower latitudes (F = 15.91, r^2^ = 0.03, Z = 3.05, p = 0.0003).

**Figure 3.**
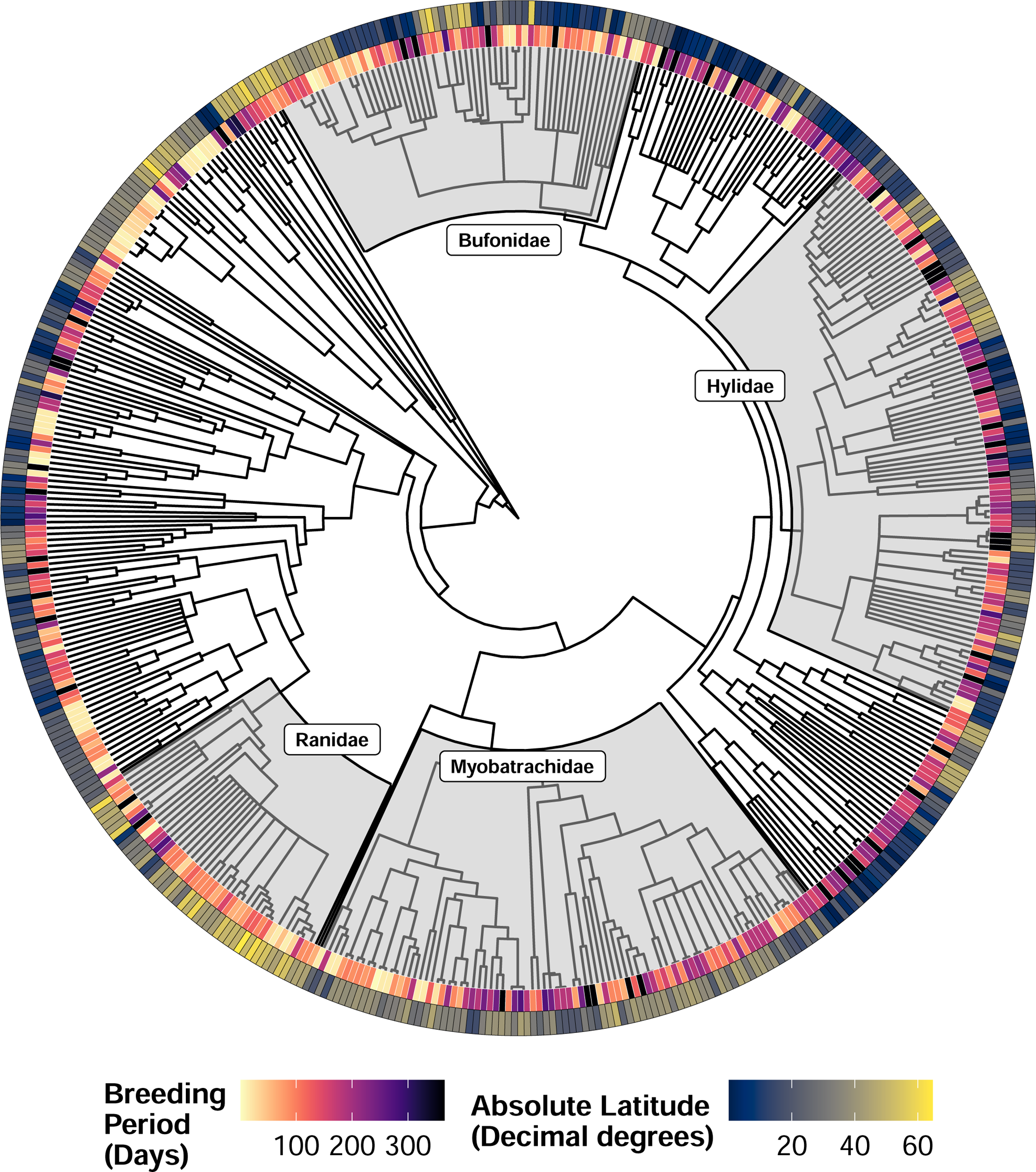
Phylogenetic differences in average breeding period and Absolute (Mean) Latitude across 497 species of frogs. Phylogeny of 497 species with breeding period and absolute mean latitude mapped at the tips. Node labels identify the 4 highest-sampled families in our dataset. Inner ring shows species means of breeding period and outer ring shows species means of absolute latitude. Anurans breed for a mean of 148 days (range = 1–365 days, std. error = 4 days). However, temperate species breed for an average of 128 days (range = 1–365 days, std. error = 5 days) and tropical species breed for an average of 174 days (range = 5–365 days, std. error = 7 days).

### Global scale analyses reveal latitude, climate, and body size drive breeding period diversity

We found breeding periods exhibited a latitudinal gradient (Fig. 4a). However, the effect of climate on breeding periods depended on body size (Fig. 4b; Fig. S1.1 in Appendix S1), as we found an interaction between climate PC1 and body size (*r^2^* = 0.07; Table 2, Rows 1, 8, 10). Relatively larger frogs exhibited longer breeding periods in the tropics (high values of PC1) defined by warm and wet environments with less temperature seasonality, compared to relatively smaller frogs which exhibited longer breeding periods in the temperate zone (low values of PC1). Notably, the geographical distribution of climate represented as scores on PC1 exhibited a strong latitudinal gradient (Fig. 4c) and this is consistent with the strong negative correlation with absolute latitude. We also determined the climate-only model (r^2^ = 0.13) was significantly improved (F = 2.88, p = 0.0003) by adding body size, microhabitat, and interactions between body size and climate (r^2^ = 0.21). Finally, we found burrowing frogs, which tend to inhabit desert microhabitats, did not exhibit relatively shorter breeding periods after correcting for multiple comparisons at the global scale (or within the temperate or tropical zones; Tables S1.1–3, Fig. S1.2).

**Figure 4.**
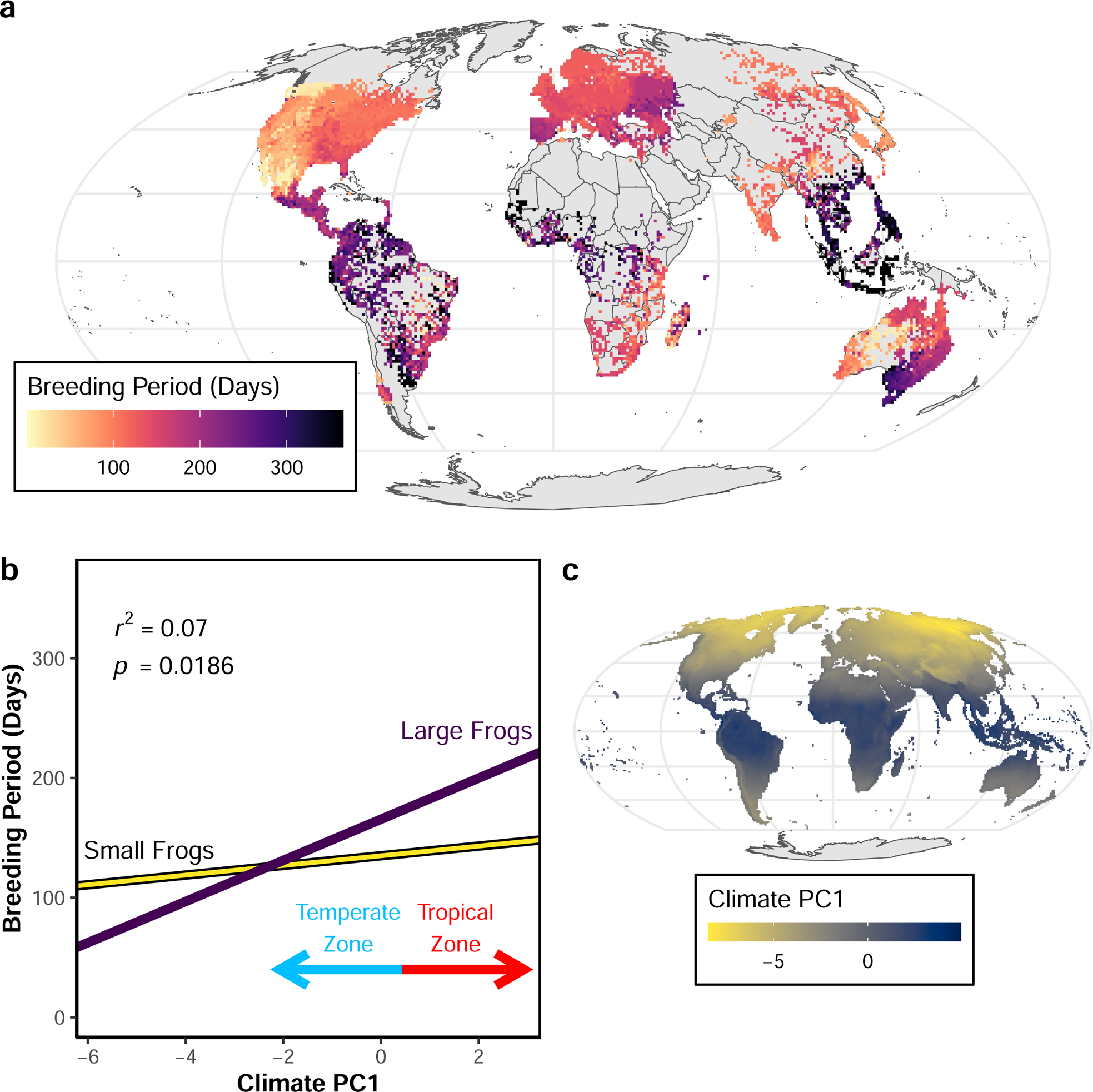
Global biogeography of breeding periods, climate PC1, and body size in 497 frog species. **a.** Mollweide equal-area projection showing pixels corresponding to 100 km^2^ areas. Pixel colors represent mean breeding periods of all species found within a pixel. **b.** The relationship between breeding period, climate PC1 (latitude), and body size. Breeding periods and PC1 scores are species means. Regression lines show the positive relationships between breeding periods and PC1 at body sizes corresponding to the 10th (24 mm) and 90th (80.48 mm) percentiles of snout-vent length after accounting for phylogenetic relationships. Larger frogs exhibit a steeper relationship between breeding periods and PC1 and larger species have longer breeding periods in the tropics while smaller species have longer breeding periods in the temperate zone. PC1 and body size explain *r^2^*= 0.07 of breeding period diversity and their interaction is significant (*p* = 0.0176). High scores on PC1 indicate tropical latitudes and low scores indicate temperate latitudes. We used a regression of absolute latitude on PC1 to determine the PC score which corresponds to 23.43628 decimal degrees. Individual points were removed for clarity but all data is displayed in Fig. S1.1. **c.** Mollweide equal-area projection showing pixels corresponding to 100 km^2^ areas. Pixel colors represent PC1 scores of climate found within a pixel.

**Table 2.** Global model summary of effects of climate and body size on breeding periods in 497 frog species. Climate shown as principal components (PC’s). Body size is snout-vent length (SVL) in mm. *Df* = degrees of freedom, *SS* = sum of squares, *MS* = mean square, *r^2^* = coefficient of determination, *F* = *F*-statistic, *Z* = standardized effect size (empirical *Z*-score), and *p* = *p*-value. Significant terms (*p* < 0.05), assessed using a consensus tree (see Methods), are in bold. Asterisks indicate significant terms supported in >50% of 1000 phylogenetic trees.

| Row | Term | Df | SS | MS | $r^2$ | *F* | *Z* | *p* |
| --- | --- | --- | --- | --- | --- | --- | --- | --- |
| 1 | **PC1\*** | **1** | **6285.50** | **6285.50** | **0.05** | **27.00** | **3.72** | **0.0001** |
| 2 | **PC2\*** | **1** | **2339.28** | **2339.28** | **0.02** | **10.05** | **2.58** | **0.0017** |
| 3 | **PC3\*** | **1** | **6025.55** | **6025.55** | **0.04** | **25.88** | **3.71** | **0.0001** |
| 4 | PC4 | 1 | 69.39 | 69.39 | 0 | 0.30 | -0.17 | 0.5722 |
| 5 | PC5 | 1 | 822.13 | 822.13 | 0.01 | 3.53 | 1.51 | 0.0614 |
| 6 | PC6 | 1 | 878.40 | 878.40 | 0.01 | 3.77 | 1.57 | 0.0537 |
| 7 | **PC7\*** | **1** | **2183.24** | **2183.24** | **0.02** | **9.38** | **2.43** | **0.0033** |
| 8 | **SVL** | **1** | **1067.75** | **1067.75** | **0.01** | **4.59** | **1.74** | **0.0362** |
| 9 | **Microhabitat\*** | **7** | **4396.42** | **628.06** | **0.03** | **2.70** | **2.25** | **0.0117** |
| 10 | **PC1:SVL\*** | **1** | **1329.36** | **1329.36** | **0.01** | **5.71** | **1.94** | **0.0176** |
| 11 | PC2:SVL* | 1 | 1709.97 | 1709.97 | 0.01 | 7.35 | 2.22 | 0.0052 |
| 12 | PC3:SVL | 1 | 1314.35 | 1314.35 | 0.01 | 5.65 | 1.96 | 0.0155 |
| 13 | PC4:SVL | 1 | 10.93 | 10.93 | 0 | 0.05 | -1.00 | 0.8231 |
| 14 | PC5:SVL | 1 | 124.85 | 124.85 | 0 | 0.54 | 0.12 | 0.4694 |
| 15 | PC6:SVL | 1 | 59.67 | 59.67 | 0 | 0.26 | -0.29 | 0.6158 |
| 16 | PC7:SVL | 1 | 59.68 | 59.68 | 0 | 0.26 | -0.28 | 0.6104 |
| 17 | Residuals | 474 | 110350.45 | 232.81 | 0.79 |  |  |  |
| 18 | Total | 496 | 139026.91 |  |  |  |  |  |

Three other key findings were independent of latitude (PC1). First, relatively larger frogs exhibited longer breeding periods in rainy environments with low topographical wetness (high values of PC2), while relatively smaller frogs had longer breeding periods in dryer environments with high topographical wetness (low values of PC2; *r^2^* = 0.04; Table 2, Rows 2, 8, 11, Fig. S1.3). Second, we found a tendency for species living in topographically wet environments with low rain seasonality (high values of PC3) to breed for longer periods (*r^2^* = 0.04; Table 2, Row 3; Fig. S1.4). Although PC3 showed an interaction with body size (Table 2, Row 12), this result was not robust to phylogenetic uncertainty (see below). Third, we found longer breeding periods were associated with warmer environments with limited potential for evapotranspiration (high values of PC7; Table 2, Row 8; Fig. S1.5). Importantly, although PC7 explained <1% of climate variance (Table 1), it explained *r^2^* = 0.02 of differences in breeding periods and had an effect size (*Z* = 2.43) comparable to other PC’s. All significant results described for this global-scale analysis were supported in 59.9–90.0% of sampled phylogenetic trees (Fig. S1.6).

### Temperate and tropical patterns are not identical to global scale patterns

We found three notable differences in the factors affecting breeding periods which differed between the global scale analysis and the temperate zone-specific analysis. First, we found a size-independent pattern where temperate species closest to the equator (high values of PC1) exhibited longer breeding periods (*r^2^* = 0.11; Table S1.4, Row 1, 10; Fig. S1.7). Second, we found that relatively larger temperate zone species exhibited longer breeding periods in topographically wet environments with low rain seasonality (high values of PC3), while relatively smaller species exhibited longer breeding periods in topographically dry environments with high rain seasonality (low values of PC3; *r^2^* = 0.08; Table S1.4, Rows 3, 8, 12; Fig. S1.8). Third, we found a size-independent pattern where temperate zone species living in rainy areas with low potential evapotranspiration (high values of PC4) exhibited longer breeding periods (*r^2^* = 0.03; Table S1.4, Row 4; Fig. 5a,b). Furthermore, as in the global scale analysis, we found a size-dependent relationship between breeding period and PC2 (*r^2^* = 0.04; Table S1.4, Rows 2, 8, 11; Fig. S1.9). Each of these results were supported in 60.6–97.4% of sampled phylogenies (Fig. S1.10). We also determined the climate-only temperate zone model (r^2^ = 0.20) was significantly improved (F = 3.87, p = 0.0001) by adding body size, microhabitat, and interactions between body size and climate (r^2^ = 0.34).

**Figure 5.**
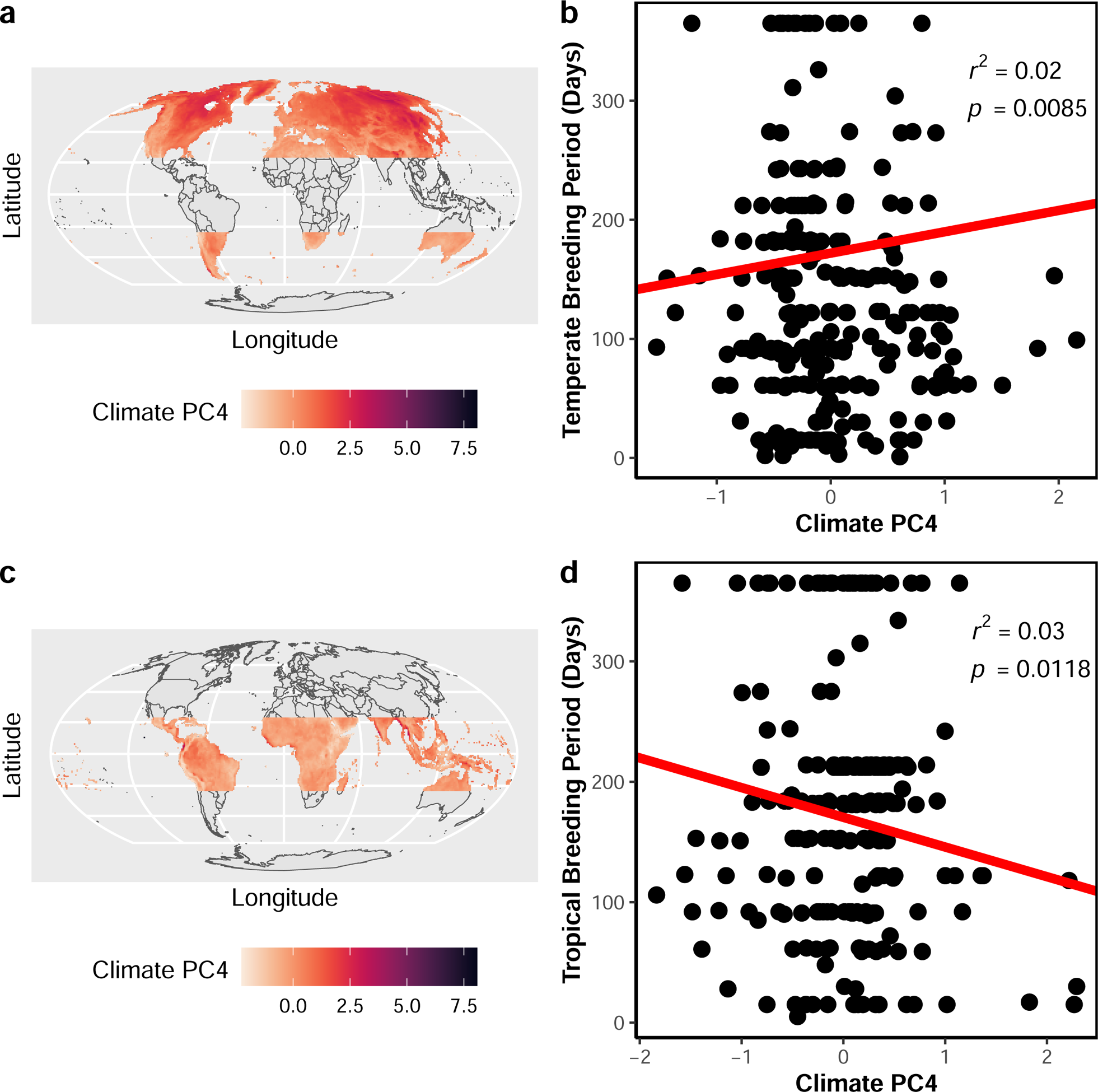
Biogeography of breeding periods and climate PC4 in the temperate and tropical zones. **a.** The marginal (positive) effect of climate PC4 on breeding periods for 279 temperate species. The red line is the regression line which accounts for phylogenetic relationships. PC4 is a contrast between annual precipitation and potential evapostranspiration; high scores are rainy areas with low potential evapotranspiration. Points are species means. **b.** Mollweide equal-area projection showing pixels corresponding to 100 km^2^ areas. Pixel colors represent PC4 scores of climate found within a pixel in the temperate zone. **c.** The marginal (negative) effect of climate PC4 on breeding periods for 218 tropical species. The red line is the regression line which accounts for phylogenetic relationships. PC4 is a contrast between annual precipitation and potential evapotranspiration; high scores are rainy areas with low potential evapotranspiration. Points are species means. **d.** Mollweide equal-area projection showing pixels corresponding to 100 km^2^ areas. Pixel colors represent PC4 scores of climate found within a pixel in the tropics.

Unlike the global and temperate zone patterns, body size did not influence any relationships between breeding period and climate in the tropical zone. As found at the global scale, we found 1) tropical species exhibited longer breeding periods in environments with low precipitation seasonality and high topographic wetness (high values of PC3; *r^2^* = 0.04; Table S1.5 Row 3; Fig. S1.11) and 2) species bred for longer periods in warmer areas with low potential for evapotranspiration (high values of PC7; *r^2^*= 0.04; Table S1.5 Row 7; Fig. S1.12). Opposite to temperate species, tropical species exhibited shorter breeding periods in rainy environments with low topographical wetness (high values of PC4; *r^2^* = 0.03; Table S1.5 Row 4; Fig. 5c,d). Each of these patterns were supported in 53.0–89.4% of sampled phylogenies (Fig. S1.13).

## Discussion

We used phylogenetic comparative methods to show how breeding periods in frogs are related to latitude, climate, and body size. We found longer breeding periods are generally associated with warmer, wetter, and less seasonal climates and body size can alter these behavior by environment relationships. However, body size and latitude were not major factors in the tropics. Overall, we found a low phylogenetic signal for breeding period and our findings support the climate and climate-size hypotheses. The zone and microhabitat hypotheses were partially and not supported, respectively. Both temperature and precipitation were important factors related to breeding periods globally, but the manner through which they impacted breeding periods differed between the temperate and tropical zones. These findings suggest that global ecology, but not phylogenetic history, is an important determinant of breeding periods in frogs.

Support for the climate-size hypothesis might be related to food availability. Warmer, wetter, and less seasonal environments may create seasonal fluxes of food resulting in environmentally constrained growth and reproduction, under the pace of life framework (Hille & Cooper, 2015). As shown previously, greater seasonality in temperature and precipitation is associated with decreased food availability (Lack, 1948; Dunn *et al*., 2000). Thus, since frogs invest more resources into reproduction when there is a greater food supply (Girish & Saidapur, 2000; Lardner & Loman, 2003), we may reasonably expect food resources to affect breeding periods. However, size-dependent (and food-independent) climate effects might explain why relatively smaller species breed for longer periods at higher latitudes if we consider the importance of energetic limitations during and after hibernation. First, frogs use metabolic energy from fat stores to overwinter and relatively smaller species have smaller fat stores (Brodeur *et al*., 2020). If smaller species begin to run low on metabolic energy during hibernation, this might result in earlier emergence from hibernation and a longer breeding period relative to larger species. Second, relatively larger species might warm slowly after emergence from hibernation such that longer and colder winters might restrict their breeding season, relative to smaller frogs. This prediction arises from the ‘heat balance hypothesis’ which describes how thermoconformers may increase heat transfer through smaller body sizes at higher latitudes (Olalla-Tarraga *et al*., 2006; Olalla-Tárraga 2011; Castro *et al*., 2021).

### Global breeding periods are composites of temperate and tropical ecology

While latitudinal gradients are typically thought of as global patterns, our study shows how global patterns of breeding period are composite patterns representing ecological dynamics that are different in temperate and tropical environments. Importantly, the global latitudinal gradient seems to reflect a latitudinal gradient in the temperate region (see Figs. 4,S1.7), as this gradient is lacking in the tropics. We also found the combined effects of body size and latitude (PC1) on global periods to be an emergent global property since we did not find this interaction within either the temperate or tropical zone. This may occur because the tropics as a whole are much warmer and wetter with less temperature seasonality compared to the temperate zone, such that differences in climate or food availability might result in unique responses of differently-sized species across latitudes and zones.

Body size and climate effects on breeding periods of tropical or temperate frogs account for global patterns. For example, we found the effects on breeding periods of regions with contrasting rain and topographic wetness (PC2) depended on body size. These effects were identical between the temperate zone and global scale. This result is consistent with the climate-size hypothesis and shows how breeding periods are determined by the combined effects of rain, topography, and body size. Additionally, longer breeding periods at the global scale resulting from less precipitation seasonality and increased topographic wetness (PC3) seem attributable to the tropics (Fig. S1.11) and large species in the temperate zone (Fig. S1.8). This effect is consistent with the climate-size hypothesis, but may also reflect the importance of body size in ephemeral water bodies used for amphibian reproduction in the temperate region. Flat, rainy, and unephemeral regions may contain fish predators which may select for slower growth rates and larger body sizes at metamorphosis due to predator-prey size asymmetry (Hecnar & M’Closkey, 1997; Lardner, 2000; Jara *et al*., 2019). We suspect predation may shorten breeding periods in smaller species, relative to larger species which benefit from less predator-prey size asymmetry. Notably, smaller frogs are predated more often in permanent water bodies (Kats *et al*., 1988; Magnusson & Hero, 1991; Summers & McKeon, 2004) and frogs reach metamorphosis slower and adulthood at a larger size in permanent water bodies filled with predators (Lardner, 2000). Finally, the global effect of warmer temperatures and diminished potential evapotranspiration (positive PC7 scores) resulting in longer breeding periods is due to the same effect in the tropics, but not in the temperate zone and is consistent with the climate hypothesis.

### Biogeographical context is crucial for understanding the drivers of breeding period

Complementing phylogenetic comparative methods with biogeographical approaches helps us understand how breeding periods (Fig. 4a) covary with climate through space. We found that frog breeding periods were shorter in the Australian Outback and longer in the southwestern Himalayas compared to regions at the same latitude. These observations coincide with dry, monsoonal regions of the northern Outback and monsoons in the southwestern Himalayas (Jeelani *et al*., 2017), given that PC2 was a contrast between precipitation and topographic wetness. These interpretations match known reproductive traits in amphibians, birds, and fishes in the Outback and Himalayas (Zann *et al*., 1995; Bentley, 1966; Joshi *et al*., 2018; Ellepola *et al*., 2021). Second, breeding periods showed a reverse-latitudinal trend in Australia and a longitudinal trend in South America and the United States. The contrast of rain seasonality and topographic wetness (PC3) in association with mountains or plains and river basins, seems to account for longer breeding periods in southern Australia, eastern United States, and in the Amazon and Pampas regions of South America. Notably, these features might explain why myobatrachid frogs (endemic to Australia and South East Asia) exhibited longer-than-expected breeding periods, relative to their average latitude. Importantly, breeding activity of frogs in northern Australia depends on heavy rains and local flooding (Tyler, 1998). Lastly, the positive effect of PC7 on global breeding periods is due to warmer temperatures and lower potential evapotranspiration. This is seemingly associated with tropical coastlines or islands and could be related to how increased conductive heat flux from exposed ground decreases evaporation in areas without canopy cover (see Lister & Garcia, 2018)

Unlike each of the previous examples, we found one variable (PC4) had opposite effects in the temperate and tropical regions. In the temperate zone, more rain and less potential evapotranspiration (high values of PC4) was conducive to longer breeding periods. This pattern is apparent in comparing the eastern and western parts of Europe, North America, and South America (e.g., the Chaco and Pampas regions). In the tropics, regions with less rain and more potential evapotranspiration led to longer breeding periods which might account for shorter than expected (relative to latitude) breeding periods in the Amazon, West Africa, southwestern India, and in the island of Borneo. These results support the climate hypothesis in the temperate zone but not in the tropical zone and might be explained if vegetative growth (shelter and breeding sites), insect growth (food), and breeding periods are restricted by too much rain and too little sunlight (low potential evapotranspiration). Typically, one might predict that less rain, not more rain, restricts plant growth. However, Huete et al. (2006) found plant growth in the Amazon rainforest is enhanced in dryer and more sunlit conditions. Thus, we infer growing seasons are longer in dryer and more sunlit environments in other tropical rainforests too, which can shorten breeding seasons in tropical frogs living in rainy environments with little sunlight. These interpretations represent hypotheses to be tested in future studies since the maps reflect community-level spatial variation while the comparative analyses summarize this variation into species means. Nonetheless, we expect both species and communities to show similar biogeographic trends relating breeding period and climate. Thus, interpreting comparative results in a biogeographic context is crucial for understanding the origins of trait variation.

### Breeding period gradients are context-dependent

In this study, we have shown how breeding periods in temperate and tropical regions are differentially impacted by climate and body size. Sometimes, the same variable may have opposite effects in each region and this may depend on how climate affects plant growth and insect abundance. This study adds to the growing body of evidence describing how the effects of environmental gradients are sometimes context-dependent (e.g., Gutiérrez-Pinto *et al*., 2014; Chaudhary *et al*., 2016; Cohen *et al*., 2018; Juarez *et al*., 2019; Martinez *et al*., 2021). While environmental gradients have historically been treated as one dimensional hypotheses, natural selection is inherently multivariate and acts on combinations of traits (see Adams & Collyer, 2019) such as breeding periods and body sizes in frogs. Longer breeding periods are associated with more egg clutches (Bull & Shine, 1979; Morrison & Hero, 2003) and larger frogs exhibit larger clutches and eggs (Prado & Haddad, 2005; Silva *et al*., 2020). Therefore, we may have found that larger frogs always exhibited longer breeding periods. Instead, our results suggest trade-offs between breeding periods and body size relative to how climate and physiology affect breeding periods across environmental gradients. Similar findings have been reported for seasonally reproducing crustaceans which show trade-offs between feeding season length and body size relative to other reproductive traits and environmental seasonality (Ejsmond *et al*., 2018).

A few other aspects of ecology seem to play an important role in determining trade-offs between breeding periods and body size. For example, shorter breeding periods in smaller species living in rainier environments with lower topographic wetness (high scores on PC2) might be related to plant diversity or predation. PC2 (Fig. S1.3) is characteristic of high-altitude rainforests (cloud forests) such as those found in Central and South America, which hold much of the world’s anuran diversity (Duellman, 1988; Giaretta *et al*., 1999; Cortés *et al*., 2008). Therefore, areas with less topographic wetness and correspondingly low vascular plant species richness (Sørensen *et al*., 2005) might limit reproductive opportunities for smaller frogs associated with leaf- or phytotelma-breeding (Donnelly & Guyer, 1994; Schulte *et al*., 2010; Poelman *et al*., 2013). Predation risk may also differ between small and large species, where larger species are less reliant on canopy cover due to their large size reducing predation pressure (Nakazawa *et al*., 2013), resulting in longer breeding periods. Alternatively, smaller frogs might face shorter breeding periods due to temporal partitioning of breeding times with other species and this depends on reproductive mode (Gottsberger & Gruber, 2004). Future research investigating the roles of plant and animal ecology on breeding periods seems likely to further uncover the roles of climate and topography in ectothermic seasonal breeders.

Further study of the relationships between body size, reproductive traits, and climate would result in a better understanding of breeding period diversity. For example, body size is related to various reproductive traits which may exhibit trade-offs themselves, including many reproductive life history traits (Furness *et al*., 2022). Additionally, previous work has found an association between decreased fecundity and warm, wet climates in terrestrial egg-laying species (Gomez-Mestre *et al*., 2012). Notably, some species may have evolved larger clutches in response to seasonal environments (Sheridan, 2009). Accounting for covariation in traits and environments is crucial for understanding the contexts and mechanisms which govern changes in seasonal reproduction.

## Data accessibility

All data and code are publicly available on datadryad.org (LINK TO BE UPDATED AFTER REVIEW).

