## Appendix 1 for "Climate and Morphology Drive Breeding Periods in Frogs"

**Data cleaning**

GBIF queries were for non-fossil and native presence coordinate observations lacking geospatial issues. We cleaned the data using the R v4.2.1 (R Core Team 2021) package *CoordinateCleaner* package v2.0-20 (Zizka *et al.* 2019) in the following ways: First, we removed non-unique coordinates for each species and data indicative of common geospatial issues (see the function *clean_coordinates* in South 2012) including 1) data georeferenced to the capital or centroid of a country, 2) data with latitudes and longitudes which were both equal or zero, 3) coordinates incorrectly associated with GBIF headquarters or natural history collections instead of the locations where the specimen was caught in the wild, and 4) specimens incorrectly reported as living in the ocean. For the latter, we used the 1° buffered land mass reference option to avoid removing coastal species from the dataset. Furthermore, we removed coordinates from the dataset with >100km uncertainty (100km uncertainty corresponds to 1° at the equator). Additionally, we restricted data for the invasive Cane Toad (*Rhinella marina*) to its naturally occurring range from southern United States to South America (AmphibiaWeb 2021) using a high resolution (1:10m) map of land boundaries available in the R package *rworldxtra* v1.01 (South 2012). Finally, we removed data outside the spatial 95% Confidence Interval for each species distribution with a minimum of 30 occurrences using the median absolute deviation (mad) method (Huber 1981) and a thinning resolution of 10 arc minutes (=0.167 decimal degrees). Thinning the data is necessary from a computational perspective when species have more than 10,000 records. This resulted in a final coordinate dataset of 848,572 observations (mean = 1,707 and range = 3–83,065 observations per species).

### Tables

**Table S1.1. Pairwise comparisons of breeding periods across microhabitats in global model.** Climate and body size differences were accounted for in this global model. N = 497 species. $p_{adj}$ is the p-value adjusted for multiple comparisons.

| **Comparison** | $p_{adj}$ |
|---|---|
| Aquatic:Arboreal | 1 |
| Aquatic:Burrowing | 0.9998 |
| Aquatic:Semi-aquatic | 1 |
| Aquatic:Semi-arboreal | 0.9999 |
| Aquatic:Semi-burrowing | 1 |
| Aquatic:Terrestrial | 1 |
| Aquatic:Torrential | 1 |
| Arboreal:Burrowing | 1 |
| Arboreal:Semi-aquatic | 1 |
| Arboreal:Semi-arboreal | 0.6004 |
| Arboreal:Semi-burrowing | 1 |
| Arboreal:Terrestrial | 0.9462 |
| Arboreal:Torrential | 0.9998 |
| Burrowing:Semi-aquatic | 0.9963 |
| Burrowing:Semi-arboreal | 0.6207 |
| Burrowing:Semi-burrowing | 0.9998 |
| Burrowing:Terrestrial | 0.4433 |
| Burrowing:Torrential | 0.9939 |
| Semi-aquatic:Semi-arboreal | 0.9928 |
| Semi-aquatic:Semi-burrowing | 1 |
| Semi-aquatic:Terrestrial | 0.9998 |
| Semi-aquatic:Torrential | 1 |
| Semi-arboreal:Semi-burrowing | 0.9998 |
| Semi-arboreal:Terrestrial | 0.9998 |
| Semi-arboreal:Torrential | 1 |
| Semi-burrowing:Terrestrial | 1 |
| Semi-burrowing:Torrential | 1 |
| Terrestrial:Torrential | 1 |

**Table S1.2. Pairwise comparisons of breeding periods across microhabitats in temperate model.** Climate and body size differences were accounted for in this global model. N = 279 species. $p_{adj}$ is the p-value adjusted for multiple comparisons.

| Comparison | $p_{adj}$ |
|---|---|
| Aquatic:Arboreal | 0.9998 |
| Aquatic:Burrowing | 1 |
| Aquatic:Semi-aquatic | 1 |

| | |
|---|---|
| Aquatic:Semi-arboreal | 1 |
| Aquatic:Semi-burrowing | 1 |
| Aquatic:Terrestrial | 1 |
| Aquatic:Torrential | 1 |
| Arboreal:Burrowing | 1 |
| Arboreal:Semi-aquatic | 1 |
| Arboreal:Semi-arboreal | 0.5148 |
| Arboreal:Semi-burrowing | 1 |
| Arboreal:Terrestrial | 0.7417 |
| Arboreal:Torrential | 0.9957 |
| Burrowing:Semi-aquatic | 1 |
| Burrowing:Semi-arboreal | 0.9904 |
| Burrowing:Semi-burrowing | 1 |
| Burrowing:Terrestrial | 0.9664 |
| Burrowing:Torrential | 1 |
| Semi-aquatic:Semi-arboreal | 0.9940 |
| Semi-aquatic:Semi-burrowing | 1 |
| Semi-aquatic:Terrestrial | 0.9749 |
| Semi-aquatic:Torrential | 1 |
| Semi-arboreal:Semi-burrowing | 0.9990 |
| Semi-arboreal:Terrestrial | 1 |
| Semi-arboreal:Torrential | 1 |
| Semi-burrowing:Terrestrial | 0.9998 |
| Semi-burrowing:Torrential | 1 |
| Terrestrial:Torrential | 1 |

**Table S1.3. Pairwise comparisons of breeding periods across microhabitats in tropical model.** Climate and body size differences were accounted for in this global model. N = 218 species. $p_{adj}$ is the p-value adjusted for multiple comparisons.

| Comparison | $p_{adj}$ |
|---|---|
| Aquatic:Arboreal | 1 |
| Aquatic:Burrowing | 0.9999 |
| Aquatic:Semi-aquatic | 0.9999 |
| Aquatic:Semi-arboreal | 1 |
| Aquatic:Semi-burrowing | 1 |
| Aquatic:Terrestrial | 1 |
| Aquatic:Torrential | 0.9999 |
| Arboreal:Burrowing | 0.9941 |
| Arboreal:Semi-aquatic | 0.6189 |
| Arboreal:Semi-arboreal | 0.9997 |
| Arboreal:Semi-burrowing | 0.9999 |
| Arboreal:Terrestrial | 0.9999 |
| Arboreal:Torrential | 0.9999 |
| Burrowing:Semi-aquatic | 0.1739 |
| Burrowing:Semi-arboreal | 0.9209 |

| | |
|---|---|
| Burrowing:Semi-burrowing | 0.9940 |
| Burrowing:Terrestrial | 0.7993 |
| Burrowing:Torrential | 1 |
| Semi-aquatic:Semi-arboreal | 0.9999 |
| Semi-aquatic:Semi-burrowing | 0.9999 |
| Semi-aquatic:Terrestrial | 0.8844 |
| Semi-aquatic:Torrential | 0.9992 |
| Semi-arboreal:Semi-burrowing | 1 |
| Semi-arboreal:Terrestrial | 0.9999 |
| Semi-arboreal:Torrential | 0.9999 |
| Semi-burrowing:Terrestrial | 1 |
| Semi-burrowing:Torrential | 0.9999 |
| Terrestrial:Torrential | 0.9999 |

**Table S1.4. Temperate zone model summary of effects of climate and body size on breeding periods in 279 frog species.** Climate shown as principal components (PC's). Body size is snout-vent length (SVL) in mm. *Df* = degrees of freedom, *SS* = sum of squares, *MS* = mean square, $r^2$ = coefficient of determination, *F* = *F*-statistic, *Z* = standardized effect size (empirical *Z*-score), and *p* = *p*-value. Significant terms ($p < 0.05$), assessed using a consensus tree (see Methods), are in bold. Asterisks indicate significant terms supported in >50% of 1000 phylogenetic trees.

| Row | Term | Df | SS | MS | $r^2$ | *F* | *Z* | *p* |
|---|---|---|---|---|---|---|---|---|
| **1** | **PC1*** | **1** | **9154.14** | **9154.14** | **0.11** | **42.28** | **3.96** | **0.0001** |
| **2** | **PC2*** | **1** | **2863.90** | **2863.90** | **0.03** | **13.23** | **2.79** | **0.0010** |
| **3** | **PC3*** | **1** | **2156.74** | **2156.74** | **0.03** | **9.96** | **2.51** | **0.0024** |
| **4** | **PC4*** | **1** | **1808.85** | **1808.85** | **0.02** | **8.35** | **2.27** | **0.0085** |
| 5 | PC5 | 1 | 362.14 | 362.14 | 0 | 1.67 | 0.91 | 0.1920 |
| 6 | PC6 | 1 | 148.42 | 148.42 | 0 | 0.69 | 0.30 | 0.3972 |
| 7 | PC7 | 1 | 41.97 | 41.97 | 0 | 0.19 | -0.41 | 0.6549 |
| 8 | SVL | 1 | 264.62 | 264.62 | 0 | 1.22 | 0.65 | 0.2740 |
| **9** | **Microhabitat*** | **7** | **4826.50** | **689.50** | **0.06** | **3.18** | **2.51** | **0.0065** |
| 10 | PC1:SVL | 1 | 807.05 | 807.05 | 0.01 | 3.73 | 1.53 | 0.0554 |
| **11** | **PC2:SVL*** | **1** | **1120.22** | **1120.22** | **0.01** | **5.17** | **1.86** | **0.0229** |
| **12** | **PC3:SVL*** | **1** | **4231.77** | **4231.77** | **0.05** | **19.55** | **3.35** | **0.0001** |
| 13 | PC4:SVL | 1 | 127.31 | 127.31 | 0 | 0.59 | 0.17 | 0.4456 |
| 14 | PC5:SVL | 1 | 159.39 | 159.39 | 0 | 0.74 | 0.31 | 0.3937 |
| 15 | PC6:SVL | 1 | 103.37 | 103.37 | 0 | 0.48 | 0.06 | 0.4978 |
| 16 | PC7:SVL | 1 | 0.06 | 0.06 | 0 | 0 | -2.13 | 0.9858 |
| 17 | Residuals | 256 | 55426.41 | 216.51 | 0.66 | | | |
| 18 | Total | 278 | 83602.86 | | | | | |

**Table S1.5. Tropical zone model summary of effects of climate and body size on breeding periods in 218 frog species.** Climate shown as principal components (PC's). Body size is snout-vent length (SVL) in mm. *Df* = degrees of freedom, *SS* = sum of squares, *MS* = mean square, $r^2$ = coefficient of determination, *F* = *F*-statistic, *Z* = standardized effect size (empirical *Z*-score), and *p* = *p*-value. Significant terms ($p < 0.05$), assessed using a consensus tree (see Methods), are in bold. Asterisks indicate significant terms supported in >50% of 1000 phylogenetic trees.

| Row | Term | Df | SS | MS | $r^2$ | *F* | *Z* | *p* |
|---|---|---|---|---|---|---|---|---|
| 1 | PC1 | 1 | 420.81 | 420.81 | 0.01 | 2.11 | 1.06 | 0.1474 |
| 2 | PC2 | 1 | 72.47 | 72.47 | 0 | 0.36 | -0.11 | 0.5549 |
| **3** | **PC3*** | **1** | **1920.49** | **1920.49** | **0.04** | **9.62** | **2.46** | **0.0024** |
| **4** | **PC4*** | **1** | **1294.41** | **1294.41** | **0.03** | **6.48** | **2.07** | **0.0118** |
| 5 | PC5 | 1 | 322.62 | 322.62 | 0.01 | 1.62 | 0.86 | 0.2021 |
| 6 | PC6 | 1 | 12.01 | 12.01 | 0 | 0.06 | -0.93 | 0.8080 |
| **7** | **PC7*** | **1** | **1815.12** | **1815.12** | **0.04** | **9.09** | **2.39** | **0.0028** |
| 8 | SVL | 1 | 639.26 | 639.26 | 0.01 | 3.20 | 1.40 | 0.0741 |
| 9 | Microhabitat | 7 | 2861.45 | 408.78 | 0.06 | 2.05 | 1.60 | 0.0537 |
| 10 | PC1:SVL | 1 | 186.94 | 186.94 | 0 | 0.94 | 0.47 | 0.3354 |
| 11 | PC2:SVL | 1 | 26.58 | 26.58 | 0 | 0.13 | -0.61 | 0.7187 |
| 12 | PC3:SVL | 1 | 160.25 | 160.25 | 0 | 0.80 | 0.38 | 0.3730 |
| 13 | PC4:SVL | 1 | 352.28 | 352.28 | 0.01 | 1.76 | 0.93 | 0.1896 |
| 14 | PC5:SVL | 1 | 356.66 | 356.66 | 0.01 | 1.79 | 0.93 | 0.1865 |
| 15 | PC6:SVL | 1 | 49.10 | 49.10 | 0 | 0.25 | -0.28 | 0.6149 |
| 16 | PC7:SVL | 1 | 30.39 | 30.39 | 0 | 0.15 | -0.52 | 0.6923 |
| 17 | Residuals | 195 | 38941.84 | 199.70 | 0.79 | | | |
| 18 | Total | 217 | 49462.69 | | | | | |

### Figures

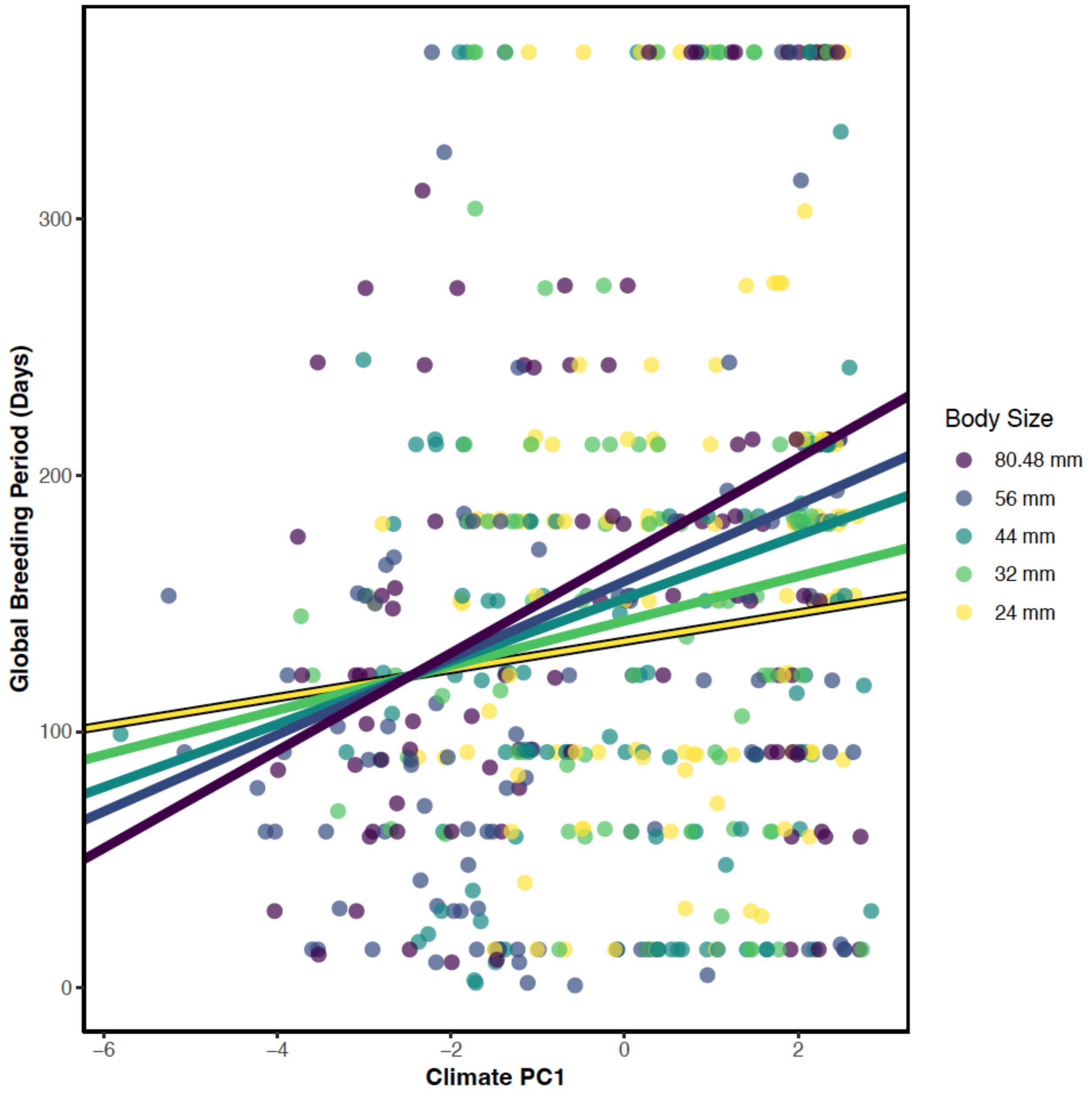


**Figure S1.1. Biogeography of breeding periods, climate PC1, and body size.** The relationship between breeding period, climate PC1 (latitude), and body size. Breeding periods and PC1 scores are species means. Regression lines show the positive relationships between breeding periods and PC1 at body sizes corresponding to the 10th (24 mm), 30th, 50th, 70th, and 90th (80.48 mm) percentiles of snout-vent length after accounting for phylogenetic relationships. Larger frogs exhibit a steeper relationship between breeding periods and PC1 and larger species have longer breeding periods in the tropics while smaller species have longer breeding periods in the temperate zone. PC1 and body size explain $r^2 = 0.07$ of breeding period diversity and their interaction is significant ($p = 0.0176$). High scores on PC1 indicate tropical latitudes and low scores indicate temperate latitudes.

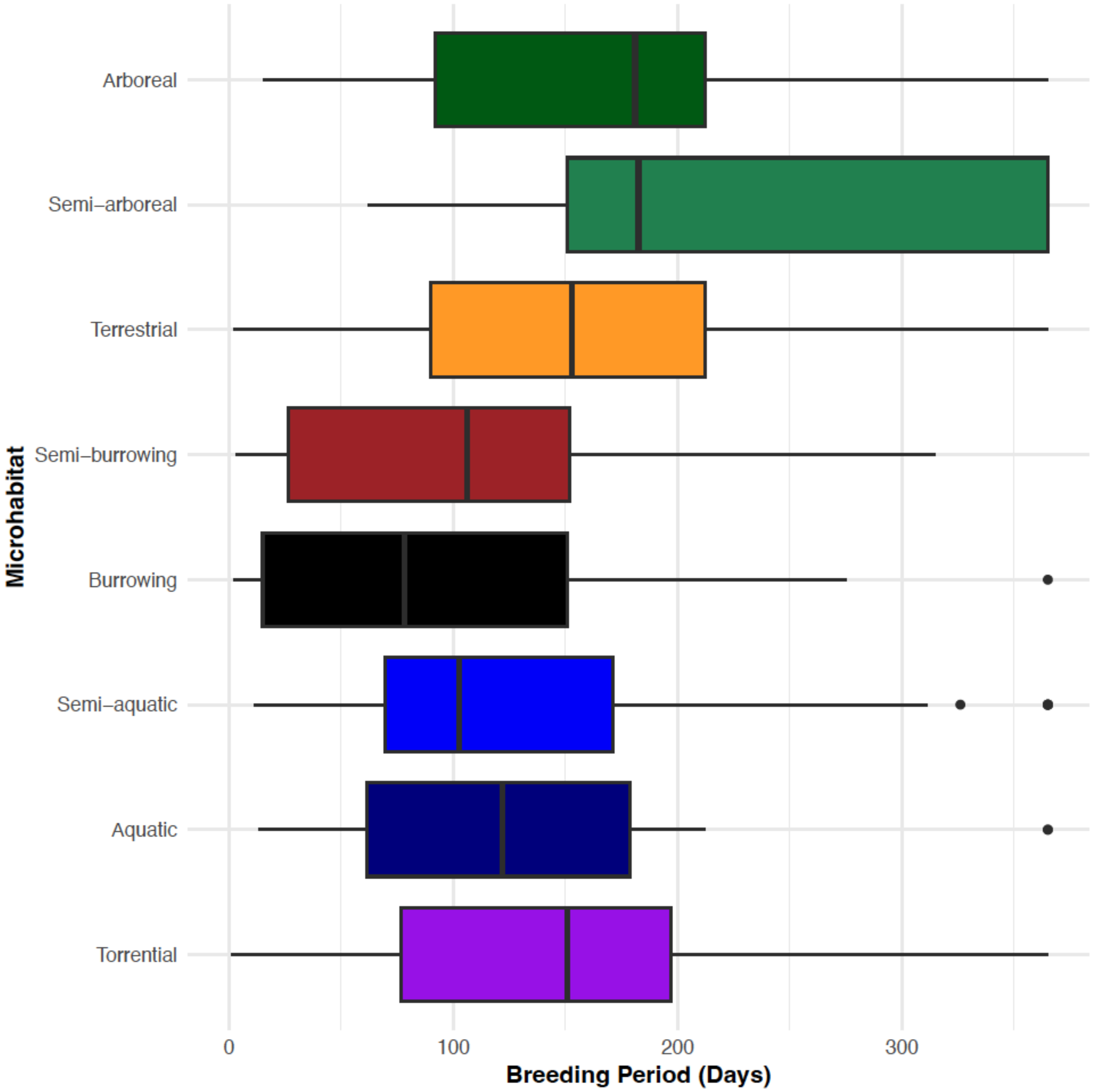


**Figure S1.2. Boxplots showing species diversity of breeding periods across microhabitats.** Microhabitats do not differ in mean breeding period after accounting for phylogenetic relatedness. Explosive breeders and year-round breeders may be found in almost all microhabitats. Species with longer breeding periods seem overrepresented in the semi-arboreal microhabitat.

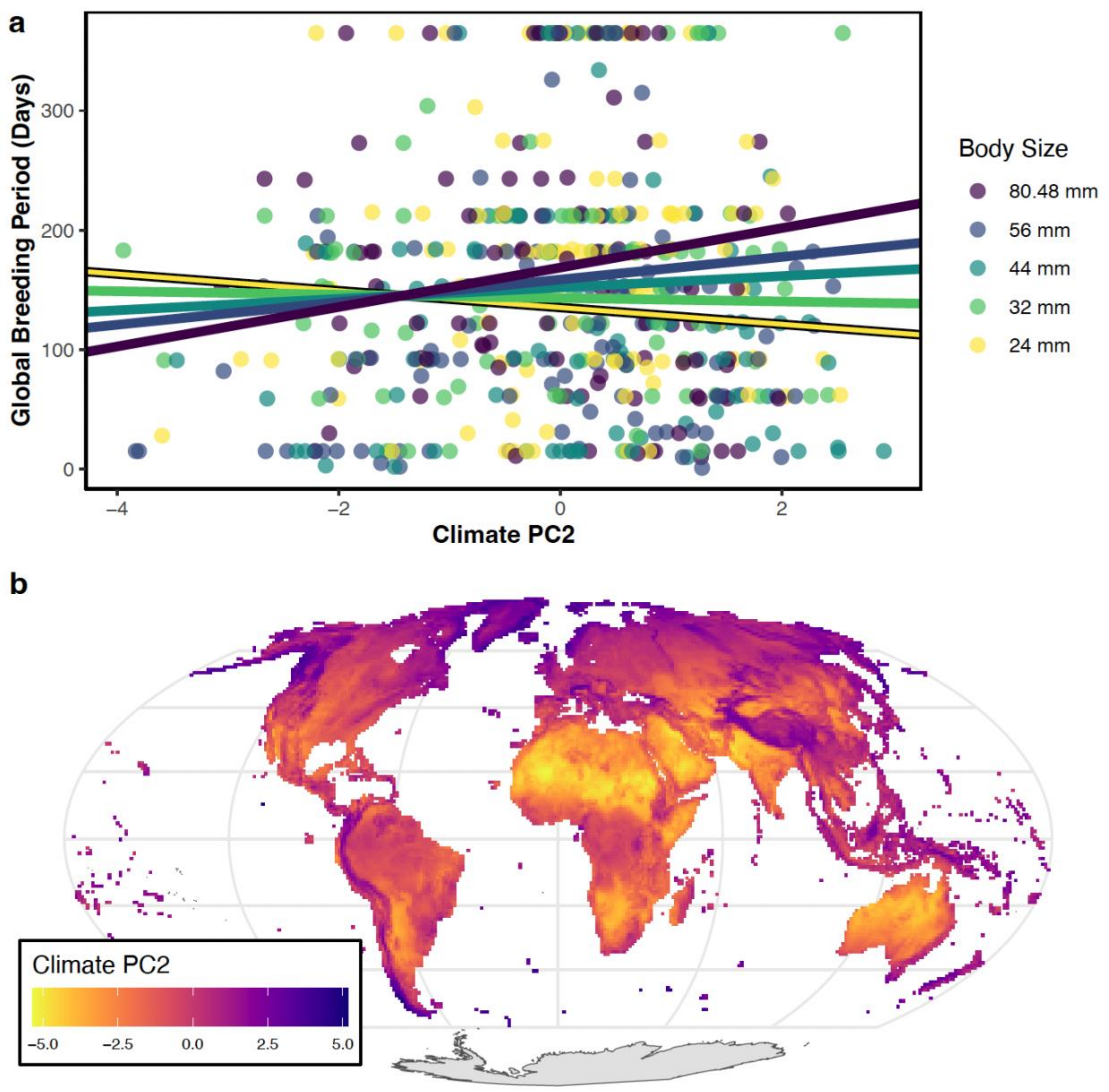


**Figure S1.3. Global biogeography of breeding periods and climate PC2**. **a.** The relationship between breeding period, climate PC2 (a contrast between annual precipitation and topographic wetness), and body size. Points are species means. High scores on PC2 indicate rainy areas with low topographic wetness. Regression lines show the positive relationships between breeding periods and PC2 at body sizes corresponding to the 10th (24 mm), 30th, 50th, 70th, and 90th (80.48 mm) percentiles of snout-vent length after accounting for phylogenetic relationships. Larger frogs exhibit a positive relationship between breeding periods and PC2 and larger species have longer breeding periods at high values of PC2 while smaller species show a negative relationship with PC2 and have longer breeding periods at low values of PC2. PC2 and body size explain $r^2$ = 0.04 of breeding period diversity and their interaction is significant ($p$ = 0.0052). **b.** Mollweide equal-area projection showing pixels corresponding to 100 $km^2$ areas. Pixel colors represent PC2 scores of climate found within a pixel.

**a**

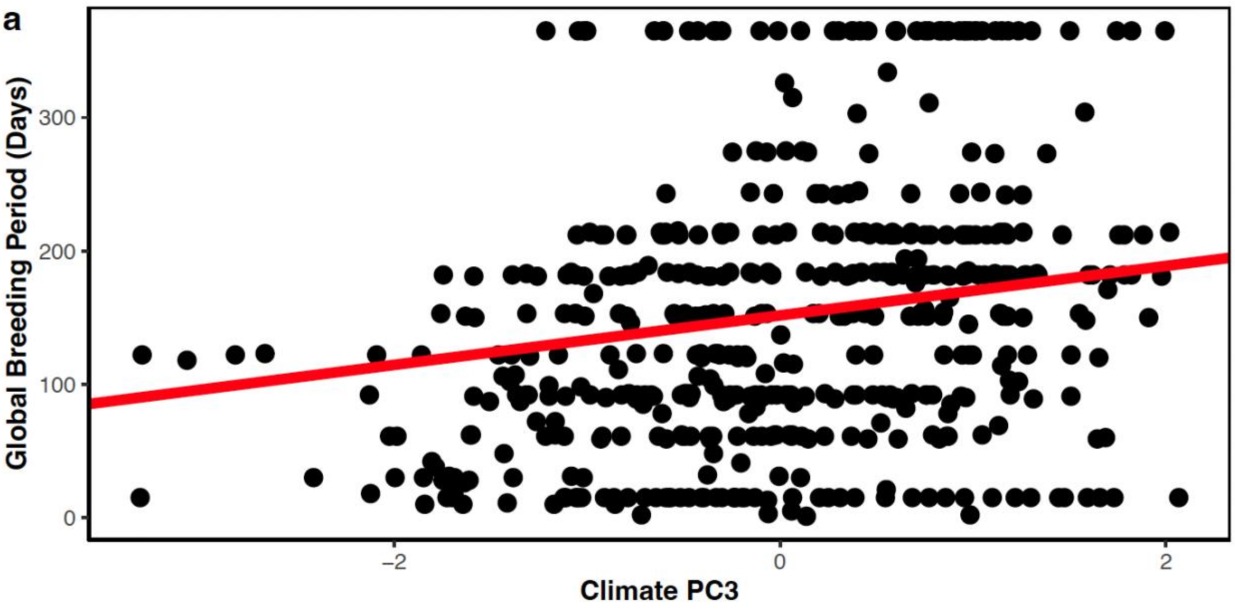


**b**

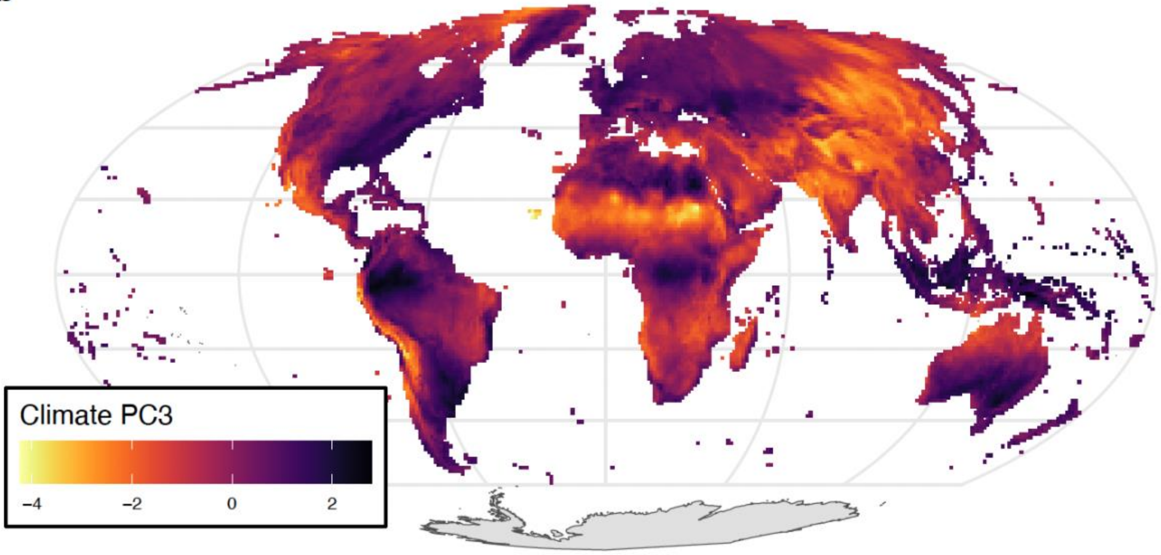


**Figure S1.4. Global biogeography of breeding periods and climate PC3**. **a.** The relationship between breeding period and climate PC3 (a contrast between topographic wetness and precipitation seasonality). Points are species means. High scores on PC3 indicate topographically wet areas with low rain seasonality. Regression line shows positive effect of PC3 on breeding periods after accounting for phylogenetic relationships. PC3 explains $r^2 = 0.04$ of breeding period diversity ($p = 0.0001$). **b.** Mollweide equal-area projection showing pixels corresponding to 100 km$^2$ areas. Pixel colors represent PC3 scores of climate found within a pixel.

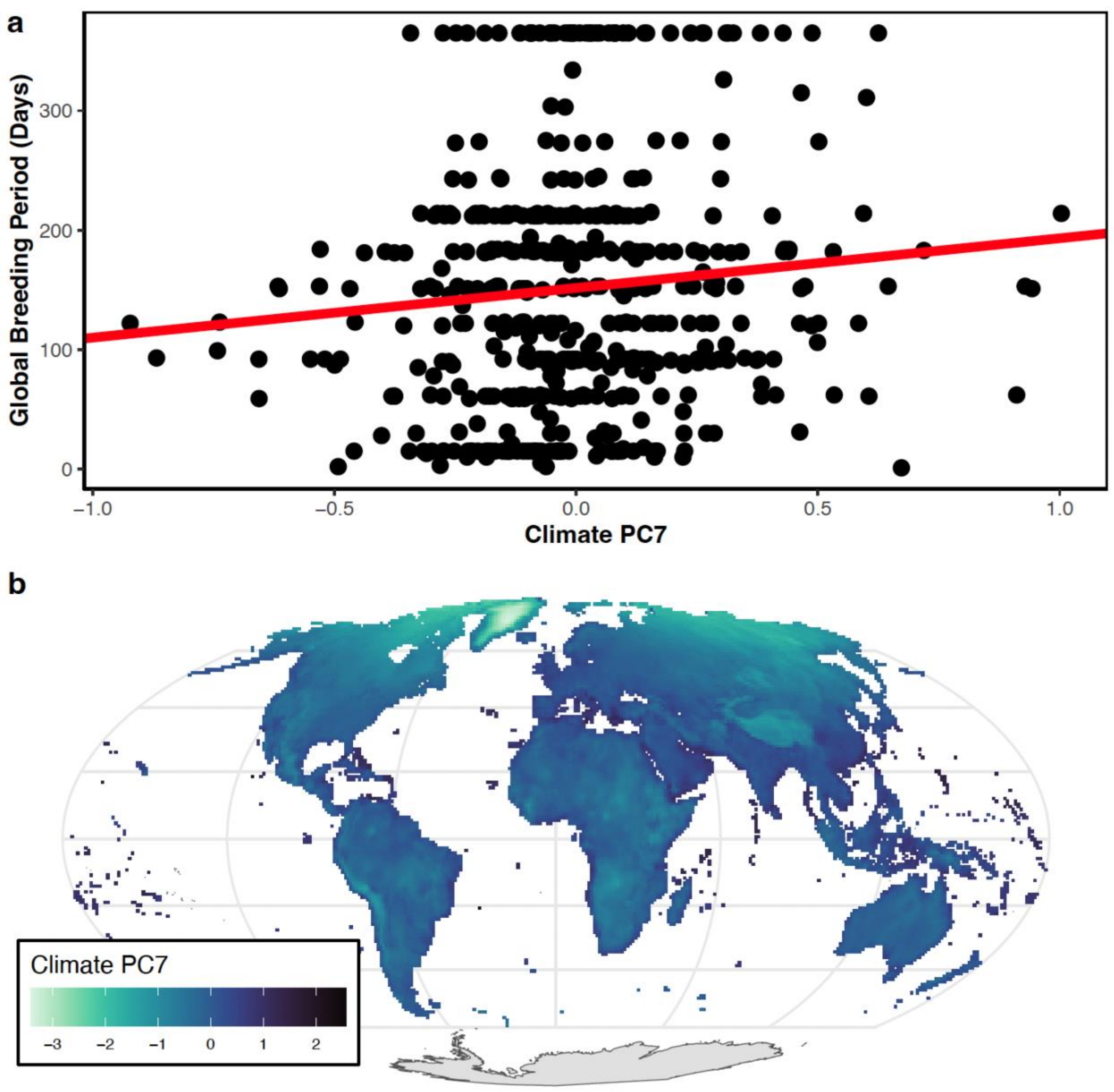


**Figure S1.5. Global biogeography of breeding periods and climate PC7**. **a.** The relationship between breeding period and climate PC7 (a contrast between annual mean temperature and annual potential evapotranspiration). Points are species means. High scores on PC7 indicate warm areas with low potential evapotranspiration. Regression line shows positive effect of PC7 on breeding periods after accounting for phylogenetic relationships. PC7 explains $r^2 = 0.02$ of breeding period diversity ($p = 0.0033$). **b.** Mollweide equal-area projection showing pixels corresponding to 100 $km^2$ areas. Pixel colors represent PC7 scores of climate found within a pixel.

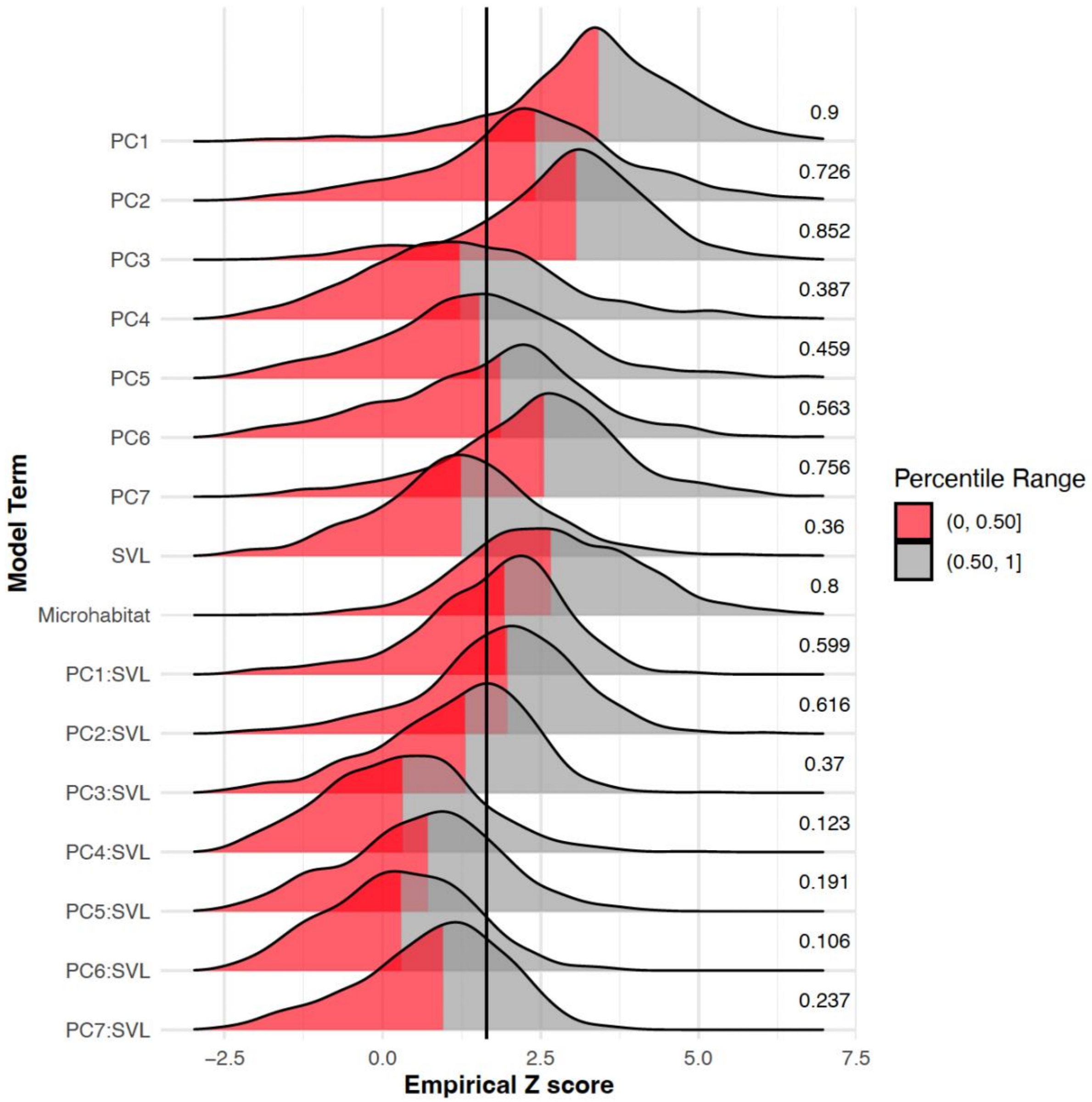


**Figure S1.6. Effect of phylogenetic uncertainty on effect sizes (z-scores) relating global climate and body size to breeding periods.** Z-scores for each model term calculated from 1,000 trees of the pseudoposterior distribution of Jetz and Pyron (2018). PC's are climate variables and SVL is snout-vent length. Vertical line denotes the $Z$ = 1.645 significance cutoff for empirically generated Z-scores. Values to the right of distributions indicate percent of significant Z-scores per distribution. Red highlights portions of the density within the 50th percentile.

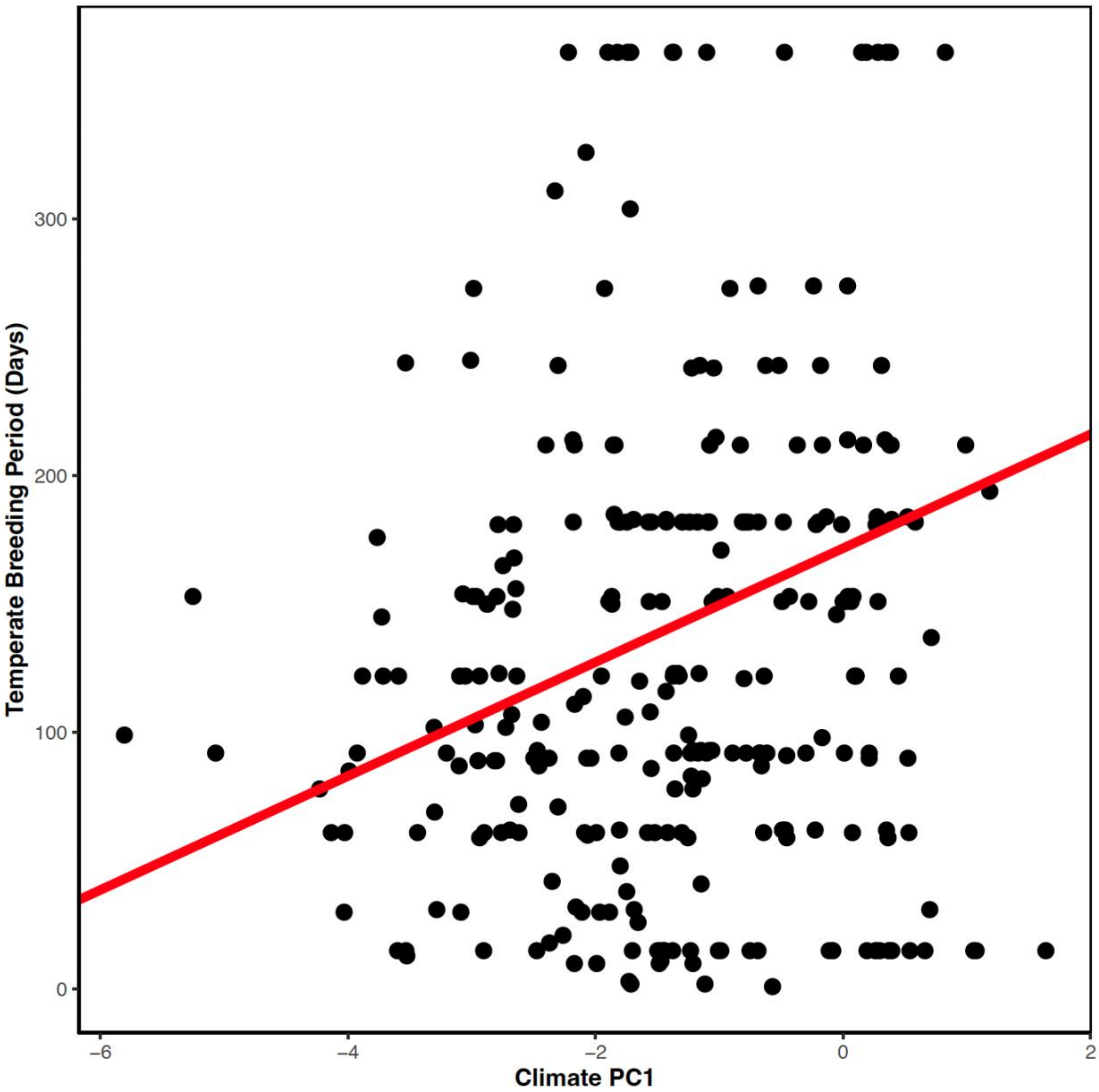


**Figure S1.7. Temperate biogeography of breeding periods and climate PC1.** The relationship between breeding period and climate PC1 (latitude). Points are species means. High scores on PC1 indicate tropical latitudes and low scores indicate temperate latitudes. Regression line shows positive effect of PC1 on breeding periods after accounting for phylogenetic relationships. PC1 explains $r^2$ = 0.11 of breeding period diversity ($p$ = 0.0001).

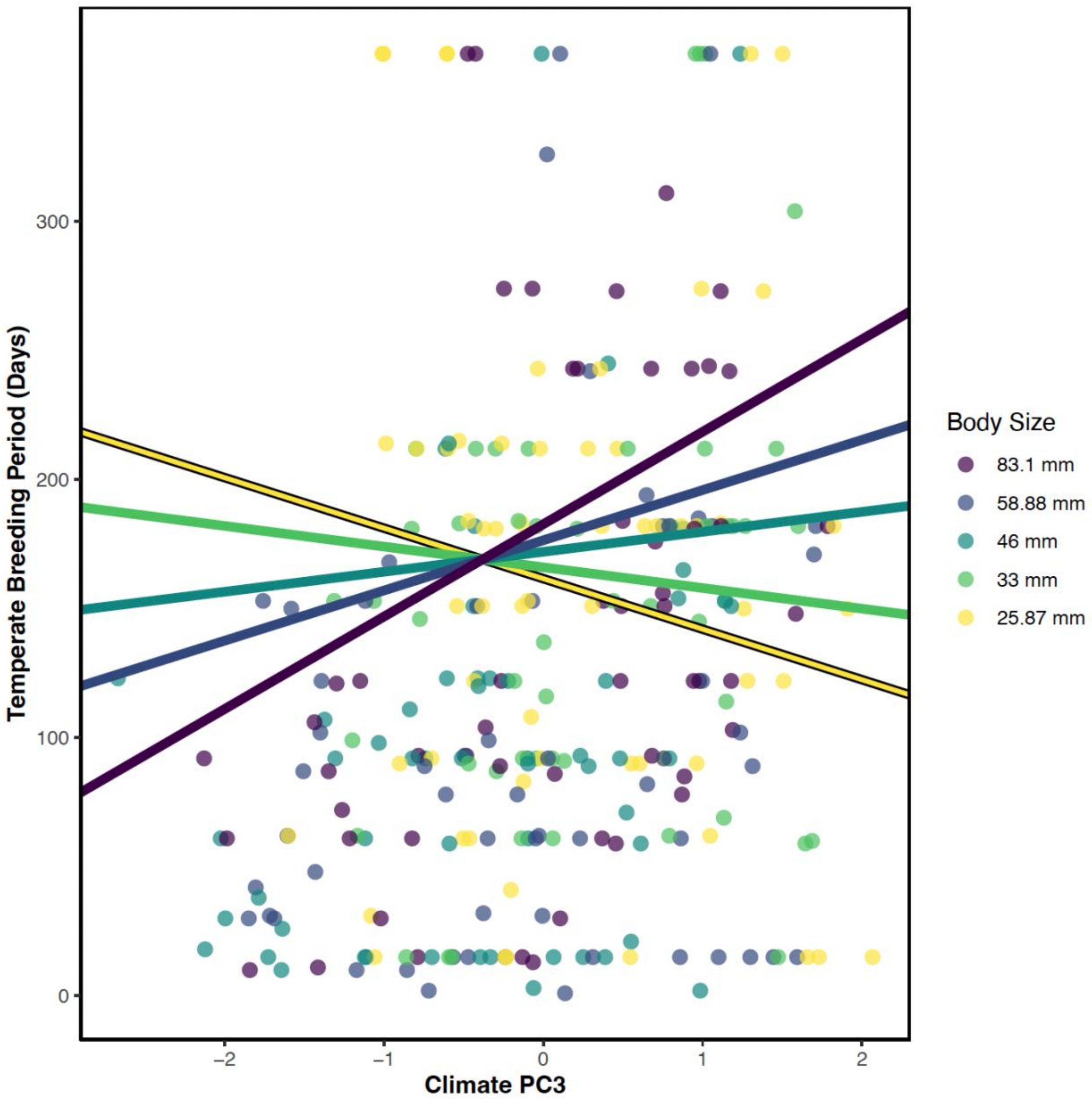


**Figure S1.8. Temperate biogeography of breeding periods and climate PC3**. The relationship between breeding period, climate PC3 (a contrast between topographic wetness and precipitation seasonality), and body size. Points are species means. High scores on PC3 indicate topographically wet areas with low rain seasonality. Regression lines show the positive relationships between breeding periods and PC3 at body sizes corresponding to the 10th (24 mm), 30th, 50th, 70th, and 90th (80.48 mm) percentiles of snout-vent length after accounting for phylogenetic relationships. Larger frogs exhibit a positive relationship between breeding periods and PC3 and larger species have longer breeding periods at high values of PC3 while smaller species show a negative relationship with PC3 and have longer breeding periods at low values of PC3. PC3 and body size explain $r^2 = 0.08$ of breeding period diversity and their interaction is significant ($p = 0.0001$).

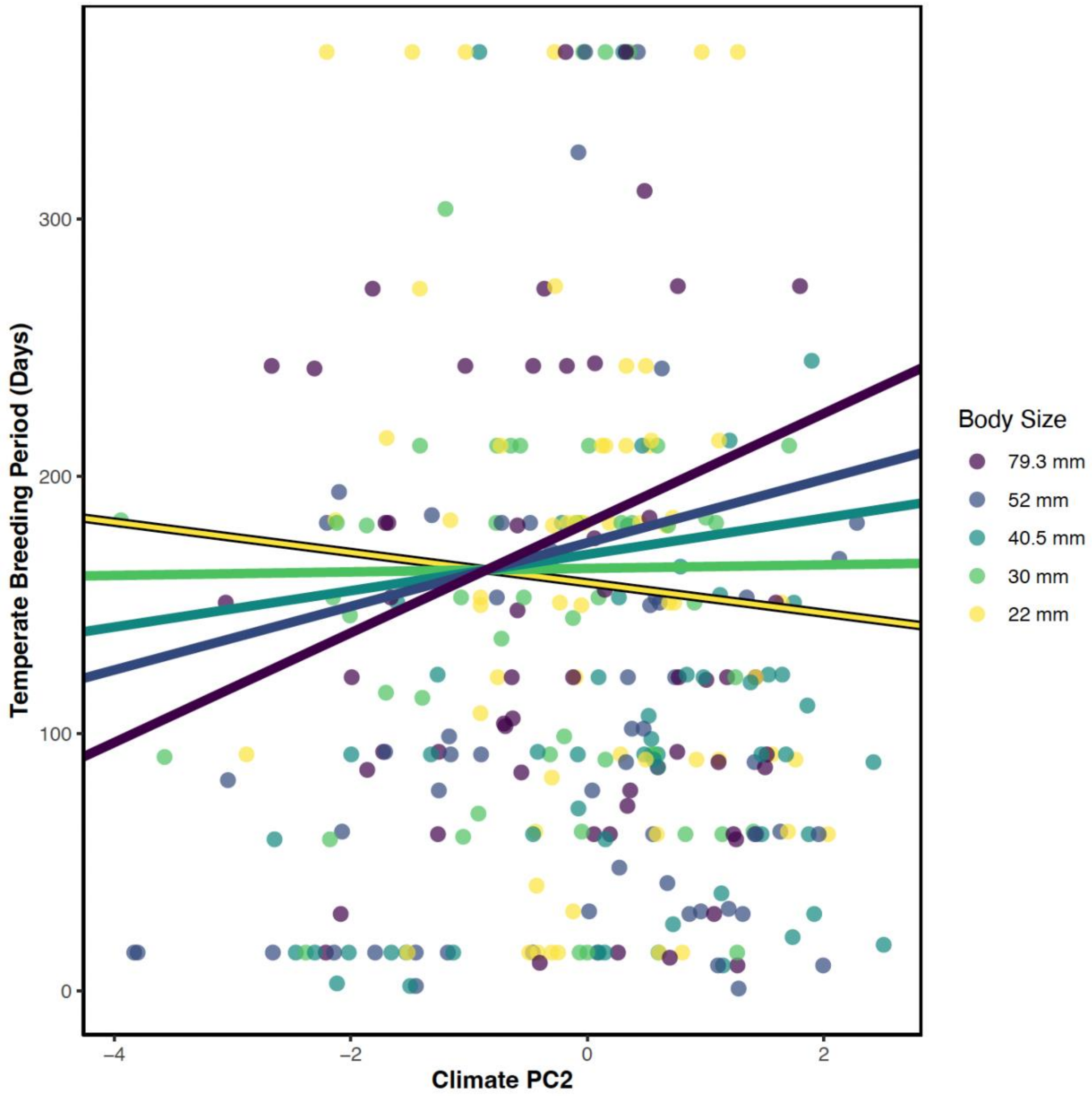


**Figure S1.9. Temperate biogeography of breeding periods and climate PC2**. The relationship between breeding period, climate PC2 (a contrast between annual precipitation and topographic wetness), and body size. Points are species means. High scores on PC2 indicate rainy areas with low topographic wetness. Regression lines show the positive relationships between breeding periods and PC2 at body sizes corresponding to the 10th (24 mm), 30th, 50th, 70th, and 90th (80.48 mm) percentiles of snout-vent length after accounting for phylogenetic relationships. Larger frogs exhibit a positive relationship between breeding periods and PC2 and larger species have longer breeding periods at high values of PC2 while smaller species show a negative relationship with PC2 and have longer breeding periods at low values of PC2. PC2 and body size explain $r^2 = 0.04$ of breeding period diversity and their interaction is significant ($p = 0.0229$).

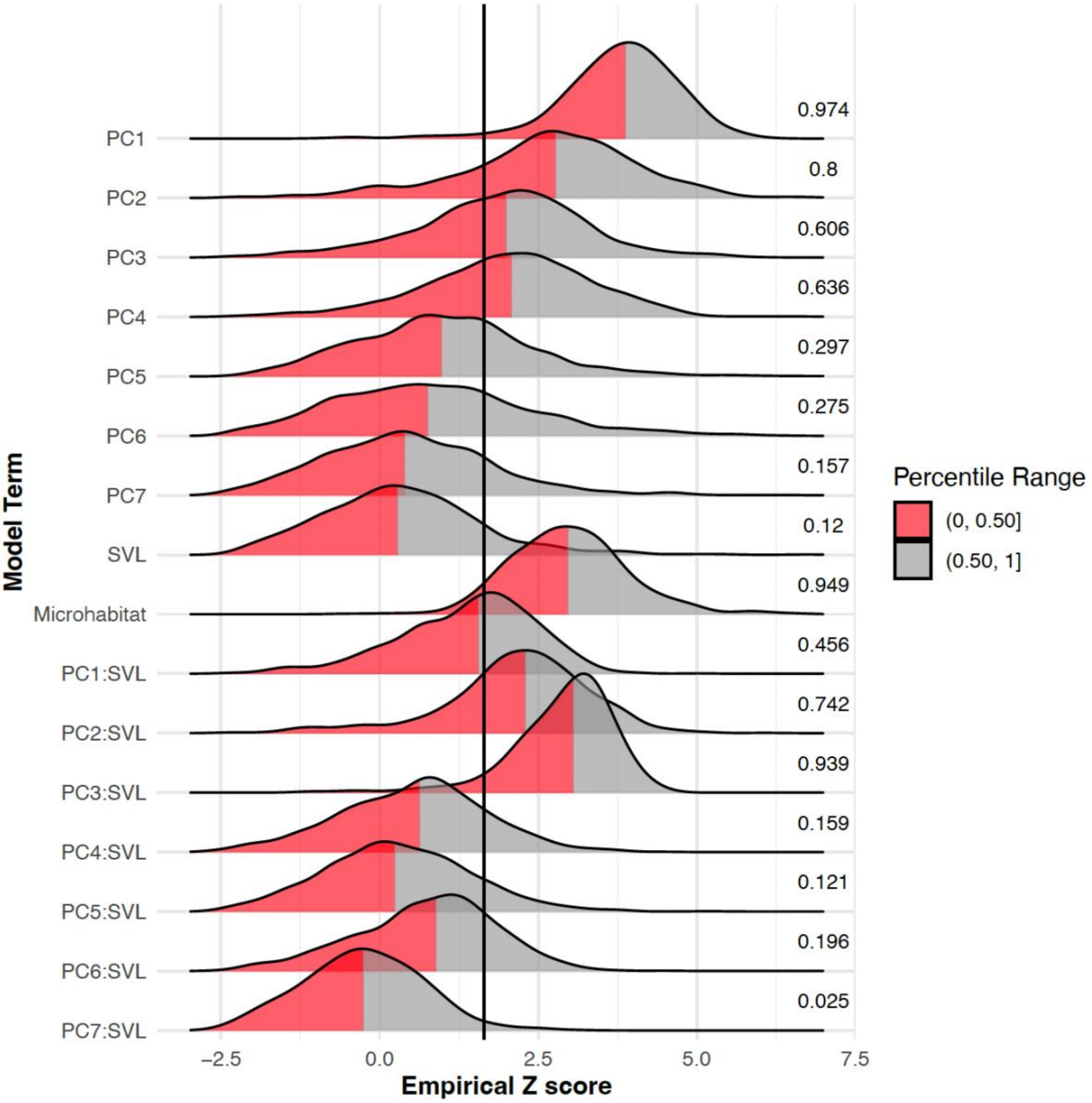


**Figure S1.10. Effect of phylogenetic uncertainty on effect sizes (z-scores) relating temperate climate and body size to breeding periods.** Z-scores for each model term calculated from 1,000 trees of the pseudoposterior distribution of Jetz and Pyron (2018). PC's are climate variables and SVL is snout-vent length. Vertical line denotes the *Z* = 1.645 significance cutoff for empirically generated Z-scores. Values to the right of distributions indicate percent of significant Z-scores per distribution. Red highlights portions of the density within the 50th percentile.

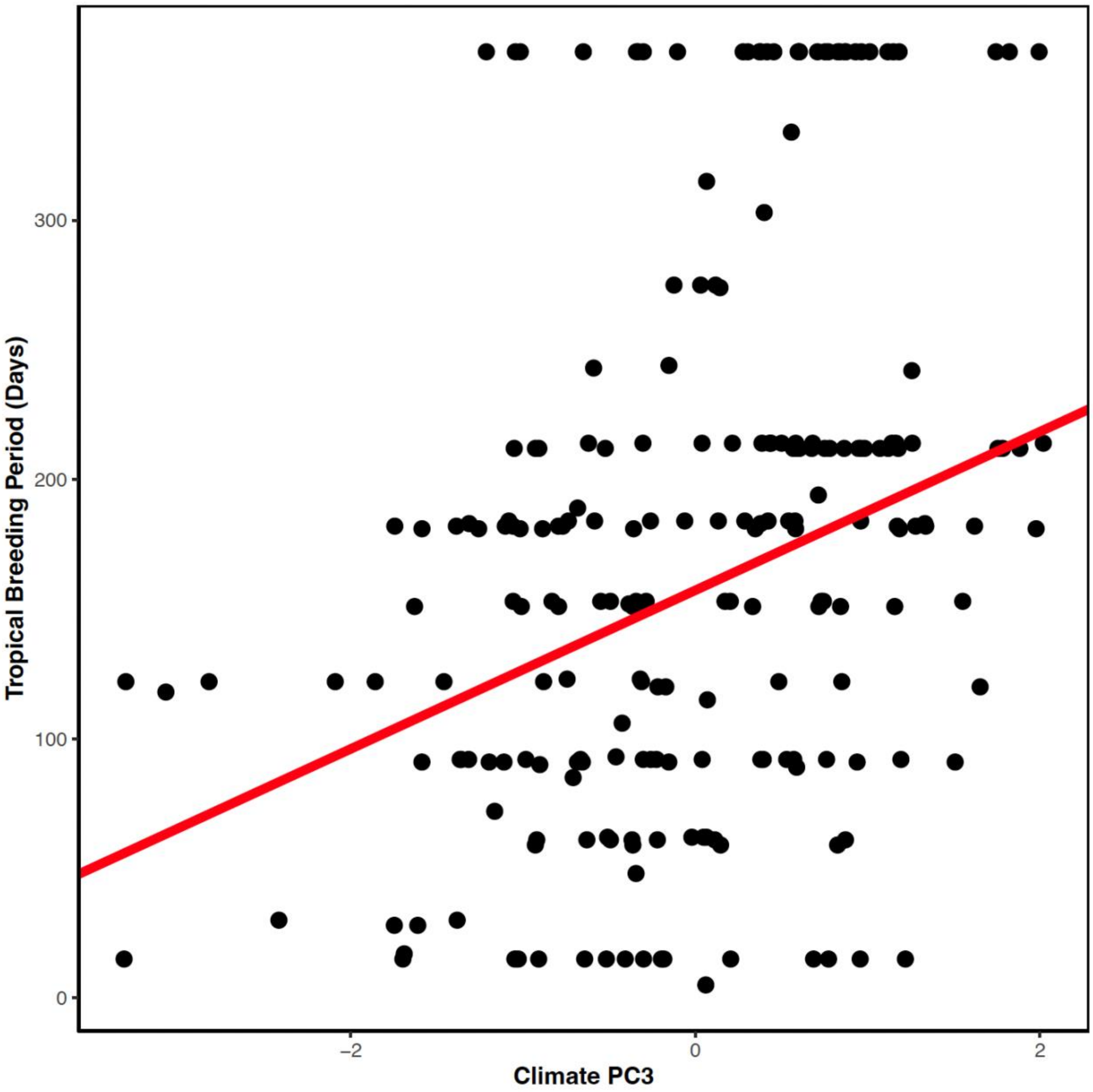


**Figure S1.11. Tropical biogeography of breeding periods and climate PC3.** The relationship between breeding period and climate PC3 (a contrast between topographic wetness and precipitation seasonality). Points are species means. High scores on PC3 indicate topographically wet areas with low rain seasonality. Regression line shows positive effect of PC3 on breeding periods after accounting for phylogenetic relationships. PC3 explains $r^2 = 0.04$ of breeding period diversity ($p = 0.0024$).

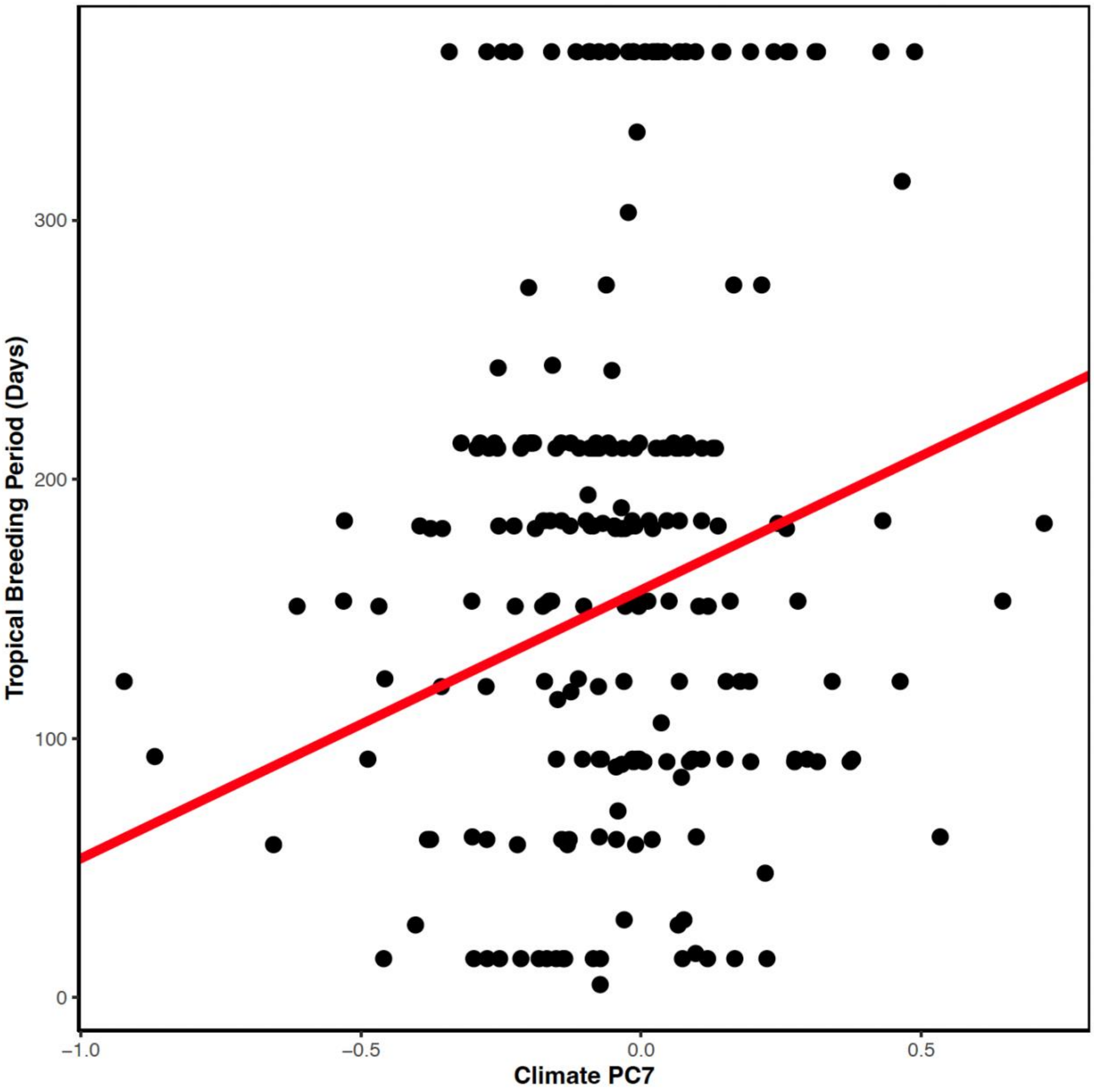


**Figure S1.12. Tropical biogeography of breeding periods and climate PC7**. The relationship between breeding period and climate PC7 (a contrast between annual mean temperature and annual potential evapotranspiration). Points are species means. High scores on PC7 indicate warm areas with low potential evapotranspiration. Regression line shows positive effect of PC7 on breeding periods after accounting for phylogenetic relationships. PC7 explains $r^2$ = 0.04 of breeding period diversity ($p$ = 0.0028).

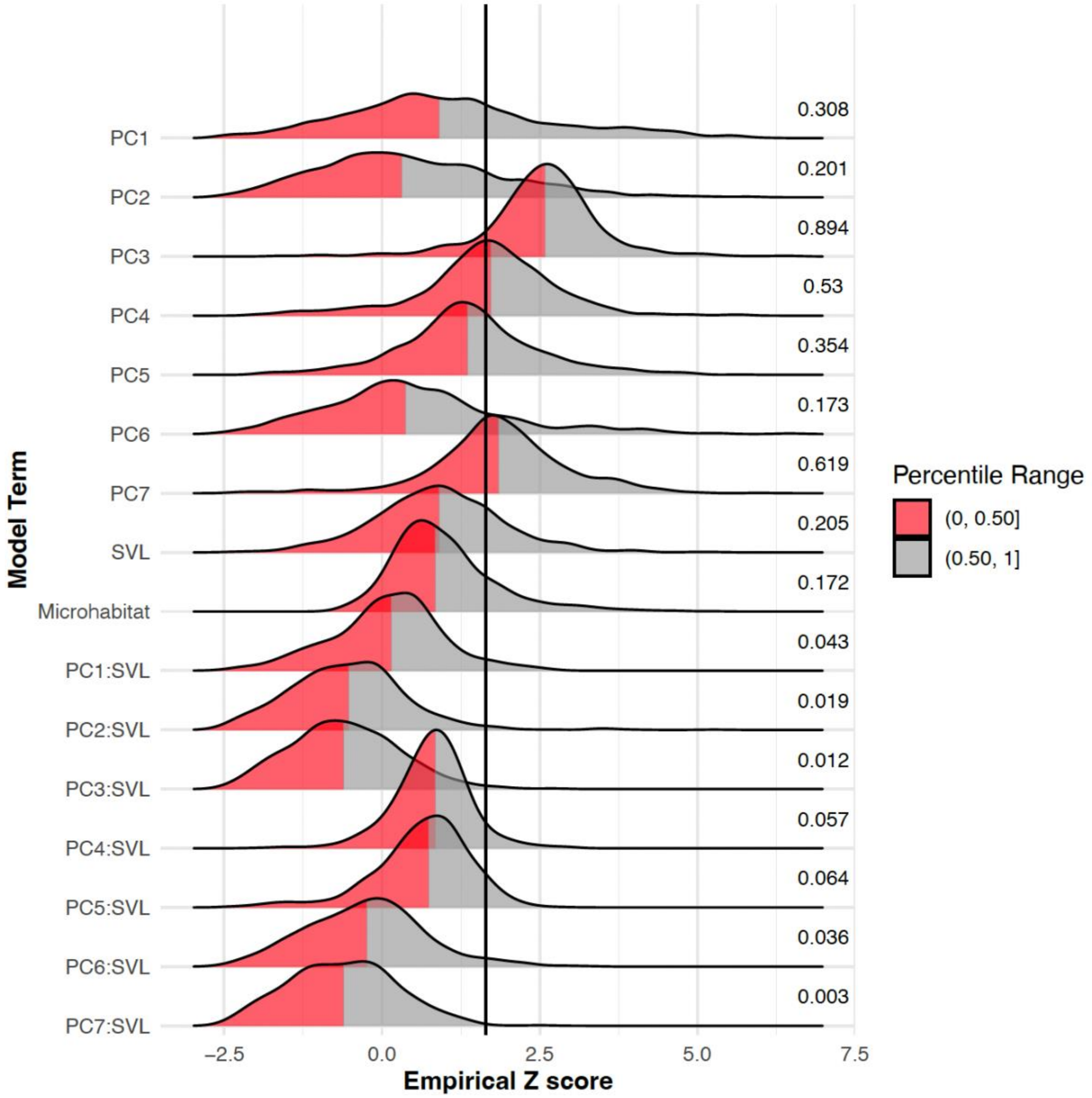


**Figure S1.13. Effect of phylogenetic uncertainty on effect sizes (z-scores) relating tropical climate and body size to breeding periods.** Z-scores for each model term calculated from 1,000 trees of the pseudoposterior distribution of Jetz and Pyron (2018). PC's are climate variables and SVL is snout-vent length. Vertical line denotes the $Z$ = 1.645 significance cutoff for empirically generated Z-scores. Values to the right of distributions indicate percent of significant Z-scores per distribution. Red highlights portions of the density within the 50th percentile.

Museum of Vertebrate Zoology, UC Berkeley. MVZ Herp Collection (Arctos). Record ID: http://arctos.database.museum/guid/MVZ:Herp:274606?seid=4060842. Source:

http://ipt.vertnet.org:8080/ipt/resource.do?r=mvz_herp (source published on 2019-07-06).

Myers, P., Espinosa, R., Parr, C.S., Jones, T., Hammond, G. S. & Dewey, T.A. (2022). *The Animal Diversity Web*. Available at: https://animaldiversity.org. Last accessed 18 July 2022.

Napoli, M. & Caramaschi, U. (2000). Description and variation of a new Brazilian species of the *Hyla rubicundula* group (Anura, Hylidae). *Alytes*, 17, 165, 184.

Ohler, A. (2007). New synonyms in specific names of frogs (Raninae) from the border regions between China, Laos and Vietnam. *Alytes*, 25, 55–74.

Ortiz, D.A., Read, M., Varela-Jaramillo, A. & Ron, S.R. (2022). *Leptodactylus petersii* In: *Anfibios del Ecuador*. Version 2021.0. { eds. Ron, S.R., Merino-Viteri, A. & Ortiz, D.A. }. Museo de Zoología, Pontificia Universidad Católica del Ecuador. Available at: https://bioweb.bio/faunaweb/amphibiaweb/FichaEspecie/Leptodactylus%20petersii. Last accessed: 17 July 2022.

Osborne, W.S. (1990). *The biology and management of the Corroboree Frog (Pseudophryne corroboree) in NSW.* Species Management Report No. 8, NPWS, Hurtsville, NSW.

Pabijan, M., Wollenberg, K.C. & Vences, M. (2012). Small body size increases the regional differentiation of populations of tropical mantellid frogs (Anura: Mantellidae). *J. Evol. Biol.*, 25, 2310–2324.

Pacheco, E.O., Ceron, K., Akieda, P.S. & Santana, D.J. (2021). Diet and morphometry of two poison frog species (Anura, Dendrobatidae) from the plateaus surrounding the Pantanal of Mato Grosso do Sul state, Brazil. *Stud. Neotrop. Fauna Environ.*, 56, 99–107.

Parker, H. W. (1940). The Australasian frogs of the family Leptodactylidae. *Novitates zoologicae*, 42, 1–106.

Peloso, P.L.V. & Sturaro, M.J. (2008). A new species of narrow-mouthed frog of the genus Chiasmocleis Méhelÿ 1904 (Anura, Microhylidae) from the Amazonian rainforest of Brazil. *Zootaxa*, 1947, 39–52.

Pengilley, R. (1973). Breeding biology of some species of Pseudophryne (Anura: Leptodactylidae) of the Southern Highlands, New South Wales. *Aust. Zool.*, 18, 15–30.

Perret, J.-L. (1966). Les amphibiens du Cameroun. *Zool. Jb. (Syst.)*, 8, 289-464.

Plăiaşu, R., Hartel, T., Băncilă, R.I., Cogălniceanu, D. & Smets, J. (2010). Comparing three body condition indices in amphibians: a case study of yellow-bellied toad *Bombina variegata*. *Amphib-reptil.*, 31, 558–562.

Poelman, E.H. & Dicke, M. (2007). Offering offspring as food to cannibals: oviposition strategies of Amazonian poison frogs (*Dendrobates ventrimaculatus*). *Evol. Ecol.*, 21, 215–227.

Pombal, J.P., and C.F.B. Haddad. 1993. *Hyla luctuosa*, a new treefrog from southeastern Brazil (Amphibia: Hylidae). Herpetologica 49: 16–21.

Pombal, J.P., Feio, R.N. & Haddad, C.F.B. (2002). A new species of torrent frog genus Hylodes (Anura: Leptodactylidae) from southeastern Brazil. *Herpetologica*, 58, 462–471.

Pombal, J.P., Sazima, I. & Haddad, C.F.B. (1994). Breeding behavior of the Pumpkin Toadlet, *Brachycephalus ephippium* (Brachycephalidae). *J. Herpetol.*, 28, 516–519.

du Preez, L. & Carruthers, V. (2009). *A complete guide to the frogs of southern Africa*. Struik Nature, Cape Town, 1–400.

R Core Team. (2022). R: A language and environment for statistical computing. R Foundation for Statistical Computing, Vienna, Austria. Available at: https://www.R-project.org/.

Radhakrishnan, C., Gopi, K.C. & Palot, M.J. (2007). Extension of range of distribution of Nasikabatrachus sahyadrensis Biju & Bossuyt (Amphibia: Anura: Nasikabatrachidae) along Western Ghats, with some insights into its bionomics. *Curr. Sci.*, 92, 213–216.

Ríos-López, N., Agosto-Torres, E., Hernández-Muñíz, R.M., Vicéns-López, C., Bernardi-Salinas, A., Tirado-Casillas, W. N., *et al.* (2015). Conservation efforts for the Puerto Rican Mountain Coqui (Anura: Eleutherodactylidae: *Eleutherodactylus portoricensis* Schmidt, 1927): Reproductive Biology in Captivity. *Life: The Excitement of Biology*, 3, 61–82.

Rodríguez-Rodríguez, E.J., Escrivà-Colomar, I. & Atiénzar, F. (2016). Reproduction of *Bufo calamita* during summer season in the Valencia province. *Bol. Asoc. Herpetol. Esp.*, 27, 51–52.

Ryan, M. (2007). Wildlife of Greater Brisbane. 3rd edition. Queensland Museum, South Brisbane 1–428.

de Sá, R.O., Grant, T., Camargo, A., Heyer, W.R., Ponssa, M.L. & Stanley, E. (2014). Systematics of the neotropical genus *Leptodactylus* Fitzinger, 1826 (Anura: Leptodactylidae): Phylogeny, the relevance of non-molecular evidence, and species accounts. *South Am. J. Herpetol.*, 9, S1–S100.

Scherz, M. (2021). Personal communication. 15 December 2021.

Señaris, J.C., Lampo, M., Rojas-Runjaic, F.J.M. & Barrio-Amorós, C.L. (2014). *Guía ilustrada de los anfibios del Parque Nacional Canaima, Venezuela*. Ediciones IVIC, Instituto Venozolano de Investigaciones

Científicas(IVIC), Caracas, Venezuela, 1–264.

Shea, G.M. & Johnston, G.R. (1988). A new species of Notaden (Anura: Leptodactylidae) from the Kimberley Division of Western Australia. *Trans. R. Soc. S. Aust.*, 112, 29–37.

Smit, G.N. (1992). Season of calling and breeding and associated control factors of a Northern Transvaal anuran population. *Afr. J. Herpetol.*, 40, 51–55.

Smith, M.J. & Roberts, J.D. (2003). Call structure may affect male mating success in the quacking frog *Crinia georgiana* (Anura: Myobatrachidae). *Behav. Ecol. Sociobiol.*, 53, 221–226.

South, A. (2012). rworldxtra: Country boundaries at high resolution. R package version 1.01. https://CRAN.R-project.org/package=rworldxtra.

Springer, L.E. & Schalk, C.M. (2016). *Lepidobatrachus laevis.* Society for the Study of Amphibians and Reptiles, 1–16. Available at: https://repositories.lib.utexas.edu/bitstream/handle/2152/44839/0904_Lepidobatrachus_laevis.pdf?sequence=1.

Stevens, R.A. (1971). A new tree-frog from Malawi. (Hyperoliinae, Amphibia). *Afr. Zool.*, 6, 313–320.

Stuart, S., Hoffmann, M., Chanson, J., Cox, N., Berridge, R., Ramani, P., *et al.* (Eds.). (2008). *Threatened amphibians of the world*. Lynx Edicions, IUCN, and Conservation International, Barcelona, Spain; Gland, Switzerland; and Arlington, Virginia, USA.

Tarkhnishvili, D.N. & Gokhelashvili, R.K. (1999). *The amphibians of the Caucasus: Advances in amphibian research in the former Soviet Union*. Pensoft Publications, Sofia, Bulgaria, 1–239.

Taylor, E.H.H. (1937). New species of hylid frogs from Mexico with comments on the rare *Hyla bistincta* Cope. *Proceedings of the Biological Society of Washington*, 50, 43–54.

Tyler, M.J. & Crook, G.A. (1987). Frogs of the Magela Creek system. *Technical Memorandum,* 19, 1–41.

Tyler, M.J., Davies, M. & Martin, A.A. (1981). New and rediscovered species of frogs from the Derby-Broome area of Western Australia. *Rec. West. Aust. Mus. Suppl.*, 9, 147–172.

Tyler, M.J. & Doughty, P. (2009). *Field guide to frogs of Western Australia*. Western Australian Museum, Perth, 1–208.

Tyler, M.J. & Knight, F. (2011). *Field guide to the frogs of Australia: Revised Edition*. Csiro Publishing, Clayton, Australia, 1–142.

Tyler, M.J., Martin, A.A. & Davies, M. (1979). Biology and systematics of a new limnodynastine genus (Anura: Leptodactylidae) from North-Western Australia. *Aust. J. Zool.*, 27, 135–150.

University of Washington Burke Museum. UWBM Herpetology Collection (Arctos). Record ID: http://arctos.database.museum/guid/UWBM:Herp:1967?seid=1894080. Source: http://ipt.vertnet.org:8080/ipt/resource.do?r=uwbm_herp (source published on 2019-07-01).

University of Washington Burke Museum. UWBM Herpetology Collection (Arctos). Record ID: http://arctos.database.museum/guid/UWBM:Herp:3342?seid=1897401. Source: http://ipt.vertnet.org:8080/ipt/resource.do?r=uwbm_herp (source published on 2019-07-01).

Verdade, V.K. & Rodrigues, M.T. (2007). Taxonomic review of Allobates (Anura, Aromobatidae) from the Atlantic Forest, Brazil. *J. Herpetol.*, 41, 566–580.

Wiens, J.J., Pyron, R.A. & Moen, D.S. (2011). Phylogenetic origins of local-scale diversity patterns and the causes of Amazonian megadiversity. *Ecol. Lett.*,14, 643–652.

Willis, Y.L., Moyle, D.L. & Baskett, T.S. (1956). Emergence, breeding, hibernation, movements and transformation of the bullfrog, *Rana catesbeiana*, in Missouri. *Copeia*, 1956, 30–41.

Wogan, G., Win, H., Thin, T., Lwin, K.S., Shein, A.K., Kyi, S. w., *et al.* (2003). A new species of Bufo (Anura: Bufonidae) from Myanmar (Burma), and redescription of the little known species *Bufo stuarti* Smith 1929. *Proceedings of the California Academy of Sciences*, 54, 141–153.

Womack, M.C. & Bell, R.C. (2020). Two-hundred million years of anuran body-size evolution in relation to geography, ecology and life history. *J. Evol. Biol.*, 33, 1417–1432.

Woodruff, D. S. (1975). Morphological and geographic variation of *Pseudophryne corroboree* (Anura: Leptodactylidae). Records of the Australian Museum, 30, 99–113.

Wells, R.W. & Wellington, C.R. 1985. A classification of the amphibia and reptilia of Australia. Australian Journal of Herpetology, Supplemental Series, 1, 1–61.

Wright, A.H. & Wright, A.A. (1949). Handbook of frogs and toads of the United States and Canada. Comstock Publishing Company, Inc., Ithaca, NY, 1–670.

Zizka A., Silvestro, D., Andermann, T., Azevedo, J., Duarte Ritter, C., Edler, D. *et al.* (2019). CoordinateCleaner: standardized cleaning of occurrence records from biological collection databases. *Methods Ecol. Evol.*, 10, 744–751. R package version 2.0-20, Available at:

https://github.com/ropensci/CoordinateCleaner. doi: 10.1111/2041-210X.13152.

Zeiner, D.C., Laudenslayer, W.F.Jr., Mayer, K.E & White, M. eds. (1988-1990). California's wildlife. Vol. I-III. California Depart. of Fish and Game, Sacramento, California. Available at: https://nrm.dfg.ca.gov/FileHandler.ashx?DocumentID=1480. Last accessed 1 October 2021.
